# MreC-MreD structure reveals a multifaceted interface that controls MreC conformation

**DOI:** 10.1101/2024.10.08.617240

**Authors:** Morgan S.A. Gilman, Irina Shlosman, Daniel D. Samé Guerra, Masy Domecillo, Elayne M. Fivenson, Claire Bourett, Thomas G. Bernhardt, Nicholas F. Polizzi, Joseph J. Loparo, Andrew C. Kruse

**Affiliations:** Department of Biological Chemistry and Molecular Pharmacology, Blavatnik Institute, Harvard Medical School, Boston, Massachusetts 02115, USA; Dana Farber Cancer Institute, Harvard Medical School Boston, Massachusetts 02115, USA; Department of Microbiology, Blavatnik Institute, Harvard Medical School, Boston, Massachusetts 02115, USA; Howard Hughes Medical Institute, Harvard Medical School, Boston, Massachusetts 02115, USA

## Abstract

The peptidoglycan (PG) cell wall is critical for bacterial growth and survival and is a primary antibiotic target. MreD is an essential accessory factor of the Rod complex, which carries out PG synthesis during elongation, yet little is known about how MreD facilitates this process. Here, we present the cryo-electron microscopy structure of *Thermus thermophilus* MreD in complex with another essential Rod complex component, MreC. The structure reveals that a periplasmic-facing pocket of MreD interacts with multiple membrane-proximal regions of MreC. We use single-molecule FRET to show that MreD controls the conformation of MreC through these contacts, inducing a state primed for Rod complex activation. Using *E. coli* as a model, we demonstrate that disrupting these interactions abolishes Rod complex activity *in vivo*. Our findings reveal the role of MreD in bacterial cell shape determination and highlight its potential as an antibiotic target.

## Introduction

The bacterial peptidoglycan (PG) cell wall is an essential structure that protects bacterial cells from osmotic lysis and is required to maintain proper cell shape. The enzymes that synthesize PG are unique to bacteria and are inhibited by the widely used β-lactam drugs. However, resistance to this class of drug now accounts for nearly 70% of antimicrobial resistance-related deaths globally^1^. Thus, new methods of blocking cell wall synthesis are needed. Regulatory pathways that direct and activate PG synthases provide an attractive alternative target for antibiotic development.

The PG heteropolymer forms a dynamic structure that must be expanded and maintained by bacterial cells during growth and division. PG is synthesized from the precursor lipid II through two enzymatic activities. Glycosyltransferases catalyze the formation of a glycan chain with alternating N-acetylglucosamine (GlcNAc) and N-acetylmuramic acid (MurNAc) sugars. Transpeptidases then crosslink the nascent glycan chain to the existing PG mesh through a peptide attached to the MurNAc sugar^2^. SEDS proteins are essential and ubiquitous glycosyltransferases that associate with class B PBP transpeptidases to carry out concerted PG synthesis during cellular elongation and division^3-10^. The specificity of SEDS-bPBPs is achieved by their association with other protein factors that form large multi-protein assemblies dedicated to PG expansion and cell division, known as the Rod complex and the divisome, respectively^11^.

The activators of the divisome complex have been well studied biochemically and structurally^12-15^. In contrast, the Rod complex has eluded structure determination and the role of its accessory factors in regulating PG synthesis remains unclear. In *E. coli*, this complex is composed of the PG synthase, RodA-PBP2, and the accessory factors MreC, MreD, and RodZ, oriented by filaments of the actin homolog MreB^16-23^. The activities of RodA and PBP2 are controlled allosterically by the structural opening of PBP2^24^. The double deletion of MreC and MreD can be rescued by mutations that stabilize the open activated state of PBP2, suggesting that these factors play a role in synthase activation^16,24,25^.

MreC is a single-pass transmembrane protein with an extended alpha-helical region that is followed by a small β-strand rich domain (β domain)^26,27^. The structure of the β domain of MreC in complex with the pedestal domain of PBP2 demonstrates that these periplasmic regions are sufficient for their interaction^28^. However, secondary structure predictions suggest that the β domain is tethered to the membrane by an extended alpha helix that could position it up to 50 Å away from its binding site on PBP2. Thus, it remains unclear how these regions access one another in the context of the full-length proteins.

Like MreC, MreD is essential in many rod-shaped bacteria. However, little is known about MreD function or its interactions with other Rod complex components. Predictions of the MreD structure suggest that it shares the same fold as the S-component proteins that serve as subunits of the energy-coupling factor (ECF) transporters used for micronutrient uptake^25,29,30^. S-components are integral membrane proteins that adopt a six transmembrane helix fold with a solvent accessible ligand binding pocket. Residues predicted to line a homologous pocket in MreD are required for its function in *E. coli*^25^, but whether MreD shares structural or functional features with the S components, and what role those features might play in Rod complex regulation, remains unknown.

To determine how MreC and MreD function in Rod complex activation, we sought to characterize the structure and function of these essential regulators. We determined the 3.6 Å resolution structure of the MreC-MreD complex, which revealed an unexpected interaction between the membrane-proximal regions of MreC and the putative ligand-binding pocket of MreD. Using single-molecule FRET, we showed that this interaction controls the conformation of MreC, stabilizing a state that is geometrically compatible with binding and allosteric activation of PBP2. Collectively, our results provide the first insights into the role of MreD in bacterial elongation and reveal a new mode of Rod complex regulation that could be targeted for drug discovery.

## Results

### Cryo-EM structure of the *Tt*MreC-MreD complex reveals three interaction interfaces

We first verified that MreC and MreD from *Thermus thermophilus* can be extracted in mild detergent and that they form a stable complex that remains associated throughout size-exclusion chromatography (Supplemental Fig. 1a,b). *Tt*MreC and *Tt*MreD form a dimer of heterodimers approximately 90 kDa in molecular weight, of which nearly 40 kDa is embedded in the detergent micelle. The small size of the complex makes it a challenging target for structure determination by cryo-electron microscopy (cryo-EM). Given this concern, we utilized a mass-enhancement strategy that relies on incorporation of an apocytochrome b562 RIL (BRIL) domain into the transmembrane (TM) helices of small membrane proteins. The recognition of the BRIL domain by an antibody fragment (Fab) increases the mass of the complex and facilitates particle alignment^31^. We additionally included a set of previously described mutations in the hinge region of the Fab heavy chain to increase its rigidity^32^. The BRIL domain was fused between transmembrane (TM) helices 4 and 5 of *Tt*MreD using a computational approach to ensure continuous incorporation of the alpha helices of the BRIL domain (see *Methods*, Supplemental Fig. 1c-f). This strategy resulted in a 3.6 Å resolution structure of the *Tt*MreC-MreD complex that resolved the transmembrane and membrane-proximal regions of both proteins (Fig. 1a,b, Supplemental Table 1, Supplemental Fig. 2a-c).

**Fig. 1.**
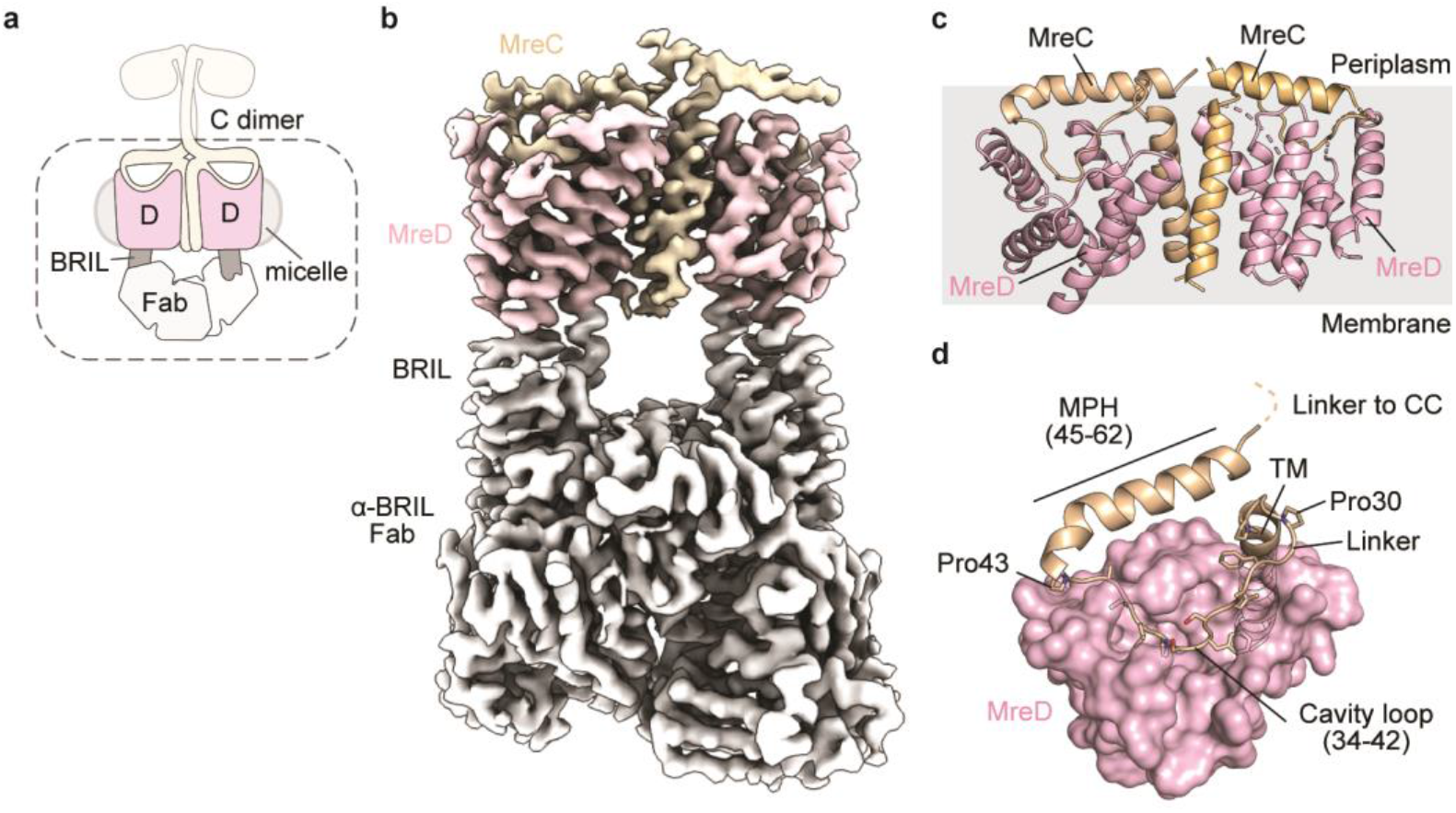
*Tt*MreC and *Tt*MreD interact through three interfaces. **a**, Schematic representation of the overall architecture of the *Tt*MreC-MreD-BRIL (rigid internal fusion) + α-BRIL Fab complex used in structure determination. The complex is shown with two MreC proteins (tan) and two MreD proteins (pink). The BRIL fusions and α-BRIL Fabs used for mass enhancement and particle alignment are shown in grey and white, respectively. The dashed line encompasses the regions that were resolved in the cryo-EM map. **b**, The cryo-EM map of the MreC and MreD complex determined to 3.6 Å resolution is shown, colored according to the cartoon in (**a**). **c**, The two copies of the transmembrane (TM) and membrane proximal regions of MreC (tan) and MreD (pink) visible in the structure are shown in ribbons. The approximate position of the membrane is annotated in grey. **d**, A single copy of MreD is shown as pink molecular surfaces. One monomer of MreC is shown as wheat ribbons. Side chains of the cavity loop and the prolines that frame this region are shown as sticks, with oxygen and nitrogen atoms colored red and blue, respectively.

*Tt*MreD interacts with the transmembrane helix of *Tt*MreC and adopts an S-component fold with a large periplasmic-facing cavity, consistent with AlphaFold predictions^25,29,33,34^ (Fig. 1c,d, Supplemental Fig. 2d). Notably, the structure revealed that the membrane-proximal regions of *Tt*MreC adopt an unexpected conformation that allows them to interact with *Tt*MreD through two additional interfaces (Fig. 1c,d). The region immediately C-terminal to the transmembrane domain of *Tt*MreC (residues 34-42) forms a loop, termed the cavity loop, that weaves into the periplasmic-facing cavity on the surface of *Tt*MreD (Fig. 1d). The cavity loop is framed by three prolines at the N- and C-termini, with an additional proline that packs against the base of the *Tt*MreD cavity. This loop is followed by an alpha helical region (residues 45-62), which we term the membrane-proximal helix (MPH). The MPH lies parallel to the membrane and contacts residues in periplasmic loop 2 and TM1 of *Tt*MreD (Supplemental Fig. 2d). Although our structure was determined in a detergent micelle, we note that the MPH has subtle amphipathic features, with charged and polar residues facing the periplasm and hydrophobic residues facing the micelle (Supplemental Fig. 2e), and thus may also interact with the bacterial inner membrane *in vivo*. The MPH is followed by a short linker region that connects to the coiled coil and the β domain, both of which were poorly resolved in our structure.

To verify that the unique architecture observed in the cryo-EM structure was not an artifact of fusing the BRIL internally to the transmembrane helices of *Tt*MreD, we designed a second fusion protein in which the BRIL domain was flexibly attached to the C-terminus. The flexibility of this fusion reduced its effectiveness in aiding particle alignment, resulting in a lower resolution structure, but removed any possible conformational constraints on the complex (Supplemental Fig. 3a). Density for the MPH was visible, oriented parallel to the membrane as was observed in the higher resolution structure. Although the cavity loop region of *Tt*MreC was poorly resolved, docking of a single copy of *Tt*MreC-MreD from the previous model agreed well with the assignment of the density in the *Tt*MreD pocket as the cavity loop (Supplemental Fig. 3b). To further validate the positioning of the cavity loop in the structure, we used a disulfide screening approach in which cysteine mutations were incorporated in both the cavity loop of *Tt*MreC and the first periplasmic loop of *Tt*MreD and tested for their ability to form a disulfide bond. For these experiments, we again used the flexible BRIL fusion construct. Two cysteine pairs formed a disulfide bond that was sensitive to redox conditions (Supplemental Fig. 3c,d). Collectively, these results verify that the membrane proximal regions of *Tt*MreC interact with *Tt*MreD.

### Structural and functional roles of the three *Tt*MreC-MreD interfaces

To determine which interfaces are required for complex formation, we designed three minimal variants of *Tt*MreC that included only its transmembrane helix and membrane-proximal regions. The first variant contained only the transmembrane helix (TM), the second contained the TM and the cavity loop, and the third contained the TM, cavity loop, and MPH. All three variants of *Tt*MreC co-purified with *Tt*MreD and remained stably associated throughout size-exclusion chromatography (Supplemental Fig. 3e,f), demonstrating that the transmembrane domains of *Tt*MreC and *Tt*MreD are sufficient for their interaction. In view of this result, we hypothesized that the role of the interaction between *Tt*MreD and the cavity loop and MPH of *Tt*MreC is not to mediate binding *per se*, but rather to functionally regulate *Tt*MreC.

To explore how *Tt*MreD binding could alter the conformation of *Tt*MreC, we first sought to assess the structure of apo-MreC. Since full-length *Tt*MreC is not amenable to structural studies in the absence of *Tt*MreD, we modeled apo-MreC using AlphaFold3 (AF3) (Supplemental Fig. 4)^34^. These models consistently predicted that the MPH forms a continuous helix with the base of the coiled coil region in *Tt*MreC, increasing the length of the alpha helical domain to approximately 100 Å. Assuming that the coiled coils are perpendicular to the membrane plane, this MPH orientation would position the β domain more than 50 Å above its binding site in PBP2^17,28^. Thus, we hypothesized that that contacts between *Tt*MreC and *Tt*MreD in the membrane proximal regions shorten the alpha helical region of *Tt*MreC, bringing the β domain in proximity to TtPBP2. We developed a single-molecule FRET (smFRET) assay to examine the conformations adopted by apo-MreC and to test the impact of MreD binding on the height of the β domain.

### MreD binding decreases the height of the MreC β domain

To measure the structural dynamics of *Tt*MreC, we constructed a monomeric version of *Tt*MreC (Fig. 2, *Methods*), since dimeric apo MreC is not amenable to smFRET measurements. We introduced cysteine substitutions to monitor the height of the β domain relative to the membrane plane: one in the β domain (*Tt*MreC^T201C^) and the others in the flexible linker that connects the MreC transmembrane helix with the cavity loop (*Tt*MreC^A33C^ or *Tt*MreC^A29C^, Fig. 2a). The resulting constructs were monodispersed on SEC and labeled specifically with sulfo-Cy3 and sulfo-Cy5 (Supplemental Fig. 5a,b). In our smFRET assay, the vertical orientation of the MPH predicted for the apo state would result in maximal elevation of the β domain and a low FRET signal (tall state, Fig. 2b). Positioning of MPH parallel to the membrane would decrease the height of the β domain and increase the FRET signal (short state, Fig. 2b). Single-molecule imaging of apo-MreC with both sets of labels showed a single major population centered at a low FRET efficiency value of 0.15-0.17 (Fig. 2c, Supplemental Table 2) and no detectable transitions to other states (Supplemental Fig. 5c). These data indicate that MreC exists in a single, tall configuration consistent with the β domain being positioned an average of ∼90 Å above the membrane.

**Fig. 2.**
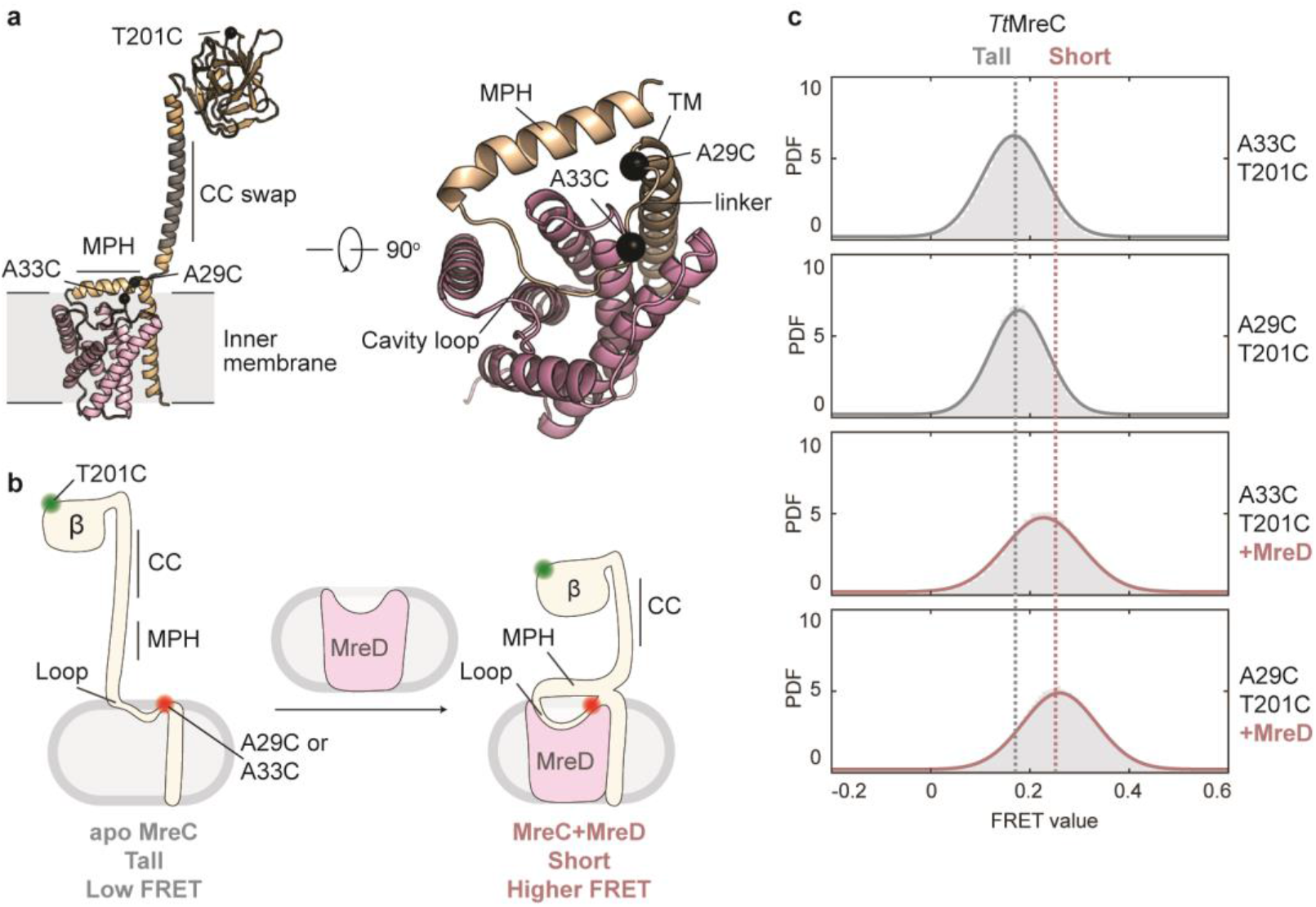
*Tt*MreD binding to *Tt*MreC changes the height of the β domain. **a**, *Tt*MreC-MreD complex with one protomer shown (left). The region of the coiled-coil (*Tt*MreC^70-96^) swapped with a monomeric helix is colored in grey. Rotated view of the *Tt*MreC-MreD membrane proximal interface (right), showing the positions of cysteine substitutions (black spheres). **b**, Schematic illustrating the smFRET assay, with one of the two possible orientations of donor (green) and acceptor (red) labels shown for simplicity. In the apo state, MreC adopts a tall state with the MPH extended, resulting in low FRET-efficiency (FE). Binding of MreD decreases the distance between the labels and increases FE. **c**, Probability density (PDF) histograms of FE values derived from single-molecule trajectories for *Tt*MreC^A33C-T201C^ and *Tt*MreC^A29C-T201C^ with and without MreD. Normal fits to the data are shown in grey (low-FRET) and pink (higher-FRET), assuming a two-state model.

Next, we tested the effect of *Tt*MreD binding on the height of the *Tt*MreC β domain by flowing excess unlabeled *Tt*MreD into the same flow cell to promote complex formation. Both *Tt*MreC cysteine constructs showed a clear shift in FRET efficiency to ∼0.25 upon addition of *Tt*MreD (Fig. 2c), consistent with a decrease in the β domain height by the length of the MPH (∼20 Å). Moreover, the *Tt*MreC-MreD complex was conformationally stable and did not exhibit any transitions to the tall state (Supplemental Fig. 5c). Our smFRET results indicate that apo-MreC adopts a tall state, but that MreD binding induces a conformational change in MreC that brings the β domain closer to the membrane, presumably through the rearrangement of the MPH.

### Membrane proximal helix and cavity loop interfaces mediate conformational changes in MreC

To determine whether MPH and cavity loop interfaces drive structural rearrangements within *Tt*MreC, we adapted our smFRET assay for the *Tt*MreC-MreD complex (*Methods*). In brief, we kept the MreC β domain label (*Tt*MreC^T201C^) and added a cysteine into MreD (*Tt*MreD^D119C^) at a position analogous to the membrane-proximal label of MreC used in the previous assay (Fig. 3a,b). Labeling of *Tt*MreC^T201C^-MreD^D119C^ (“WT”) was specific and did not disrupt complex formation (Supplemental Fig. 6a,b). Single-molecule imaging of *Tt*MreC-MreD “WT” showed a single population centered at a FRET efficiency value of ∼0.25 and no detectable structural dynamics, confirming that the complex exists in a stable short state (Fig. 3c, Supplemental Fig. 6c, Supplemental Table 2). We then designed mutations predicted to disrupt either MPH or cavity loop contacts and evaluated their conformational ensembles.

**Fig. 3.**
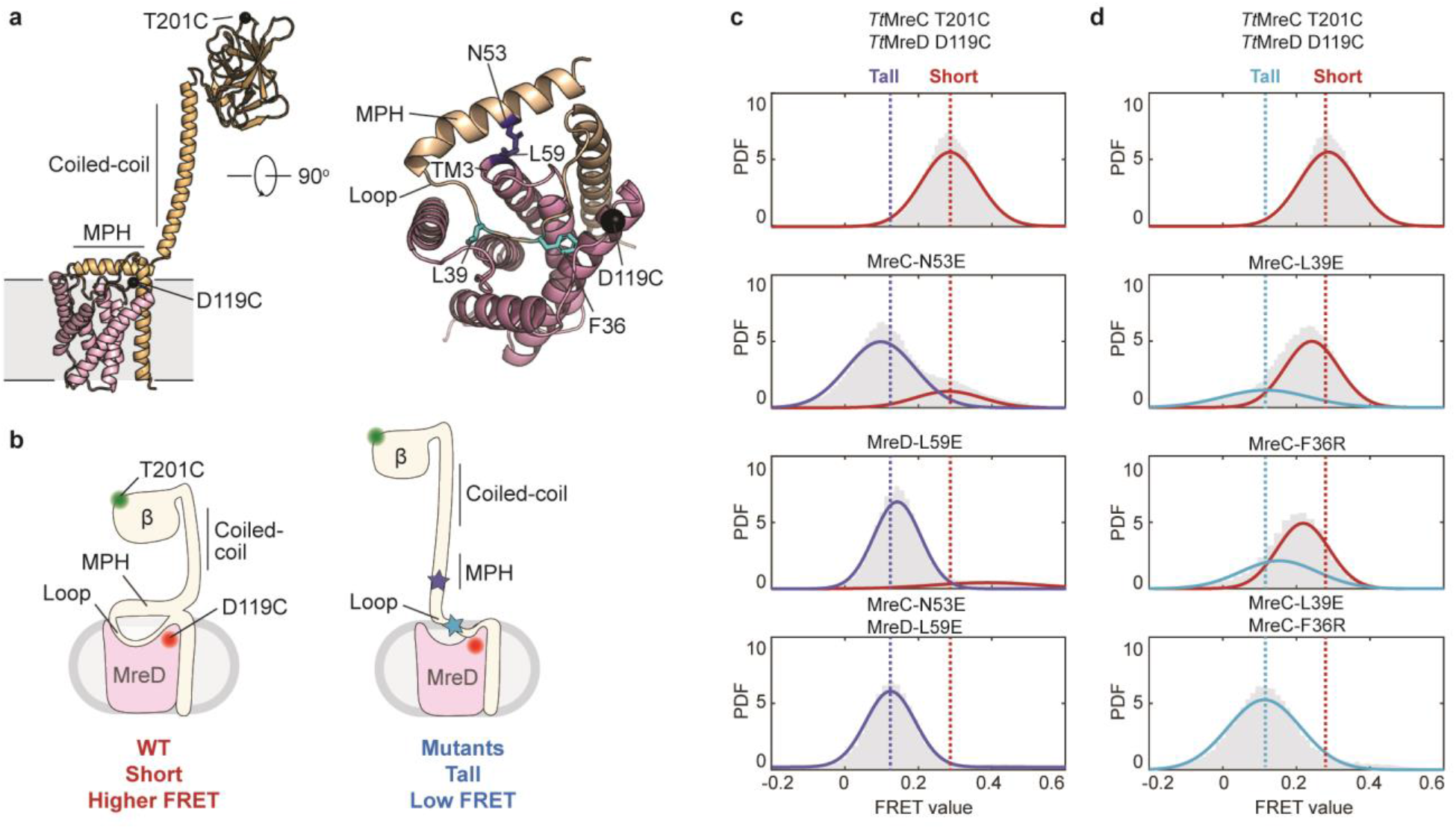
MPH and cavity loop contacts drive conformational changes in *Tt*MreC. **a**, *Tt*MreCD complex as in Fig. 1 showing the positions of cysteine substitutions as black spheres (left). Rotated view of the complex (right), showing mutations at the MPH (dark blue) and cavity loop (light blue) interfaces as sticks. **b**, Schematic illustrating the smFRET assay. WT complex adopts a short state (higher-FRET); mutations to MPH and loop interfaces induce a conformational change to a taller state (low-FRET). **c**, Probability density (PDF) histograms of FRET-efficiency (FE) values derived from single-molecule trajectories for *Tt*MreCD WT and MPH mutants. Normal fits to the data are shown in blue (low-FRET) and red (higher-FRET), assuming a two-state model. **d**, Same as (**c**) for loop mutants. *Tt*MreCD WT plot is reproduced from (**c**) for convenience.

The interface between the membrane proximal helix of *Tt*MreC and the third transmembrane helix of *Tt*MreD (TM3) is composed of only a handful of residues and has few specific interactions (Fig. 3a, right). To disrupt this interface, we substituted two residues that come in close contact with each other (*Tt*MreC^N53E^, *Tt*MreD^L59E^) with glutamate. Notably, *Tt*MreD^L59^ is at the tip of TM3 and maps to the same region as a previously identified dominant-negative mutation in *E. coli* MreD (I69F)^25^. Both mutants were biochemically well-behaved (Supplemental Fig. 6 a,b) and exhibited markedly shifted smFRET profiles relative to “WT”, with a high occupancy tall state in addition to a low occupancy short state (Fig. 3c, Supplemental Fig. 6c). When combined, the mutations had an additive effect, fully shifting the ensemble to the tall state (Fig. 3c). These results are in agreement with our model that contacts between MreC and MreD at the MPH interface drive conformational changes that bring the β domain closer to the membrane. Finally, we note that the amphipathic nature of the MPH (Supplemental Fig. 2e) likely contributes to its propensity to partition to the membrane and energetically favors the conformational transition of MreC to the short state.

To test whether cavity loop contacts contribute to the conformational changes of MPH, we destabilized the loop interface by swapping two large hydrophobic residues facing the cavity for charged residues (*Tt*MreC^L39E^, *Tt*MreC^F36R^) (Fig. 3a). Each substitution preserved binding between *Tt*MreC and *Tt*MreD (Supplemental Fig. 7a,b) and partially shifted the conformation of MreC to the tall state (Fig. 3d). The mutations also appeared to induce structural dynamics within *Tt*MreC, leading to rapid exchange between tall and short states (Supplemental Fig. 7c). We hypothesize that this motion corresponds to the wobbling of the MPH between vertical and horizontal orientations. However, because the FRET efficiency values of the two states are so close, the kinetics of this exchange could not be quantified with confidence. The double mutant showed complete loss of the short state and no detectable dynamics (Fig. 3d), suggesting that cavity loop contacts allosterically control the repositioning of the MPH and as a result, the height of the β domain. Collectively, our results demonstrate that MreC can adopt two distinct configurations, with its β domain either raised or lowered relative to the membrane plane, and that binding of MreD through the MPH and cavity loop interfaces stabilizes the short state of MreC.

The changes in height observed in the smFRET experiments are consistent with a lowering of the *Tt*MreC β domain by approximately 20 Å. However, if the coiled coil remains vertical, this would still position the β domain ∼30 Å from PBP2. How then does *Tt*MreC span the remaining distance? Although our structural studies were not able to resolve the periplasmic regions of *Tt*MreC, increasing the threshold of the cryo-EM maps allowed us to determine the approximate orientation of the coiled coil relative to the membrane plane. In the low-resolution structure, the blurred density of the coiled coil is consistent with a perpendicular but highly mobile conformation of this region (Fig. 4a-c). In contrast, the higher resolution structure captures a stably tilted state of the coiled coil that would be expected to decrease the relative height of the β domain (Fig. 4d-f). To determine if this degree of tilting is sufficient to facilitate *Tt*MreC binding to TtPBP2, we combined AF3 predictions with our experimental structure to generate a model of the four-component complex (*Tt*MreC-MreD-RodA-PBP2) (see *Methods*). The resulting model predicts that the β domain of *Tt*MreC can reach the pedestal domain of *Tt*PBP2 when the coiled coil adopts the tilted state (Supplemental Fig. 8). Thus, we hypothesize that *in vivo* the coiled-coil of MreC-MreD rapidly samples a continuum of tilted states, until, when bound to PBP2, it is stabilized in the maximally tilted conformation.

**Fig. 4.**
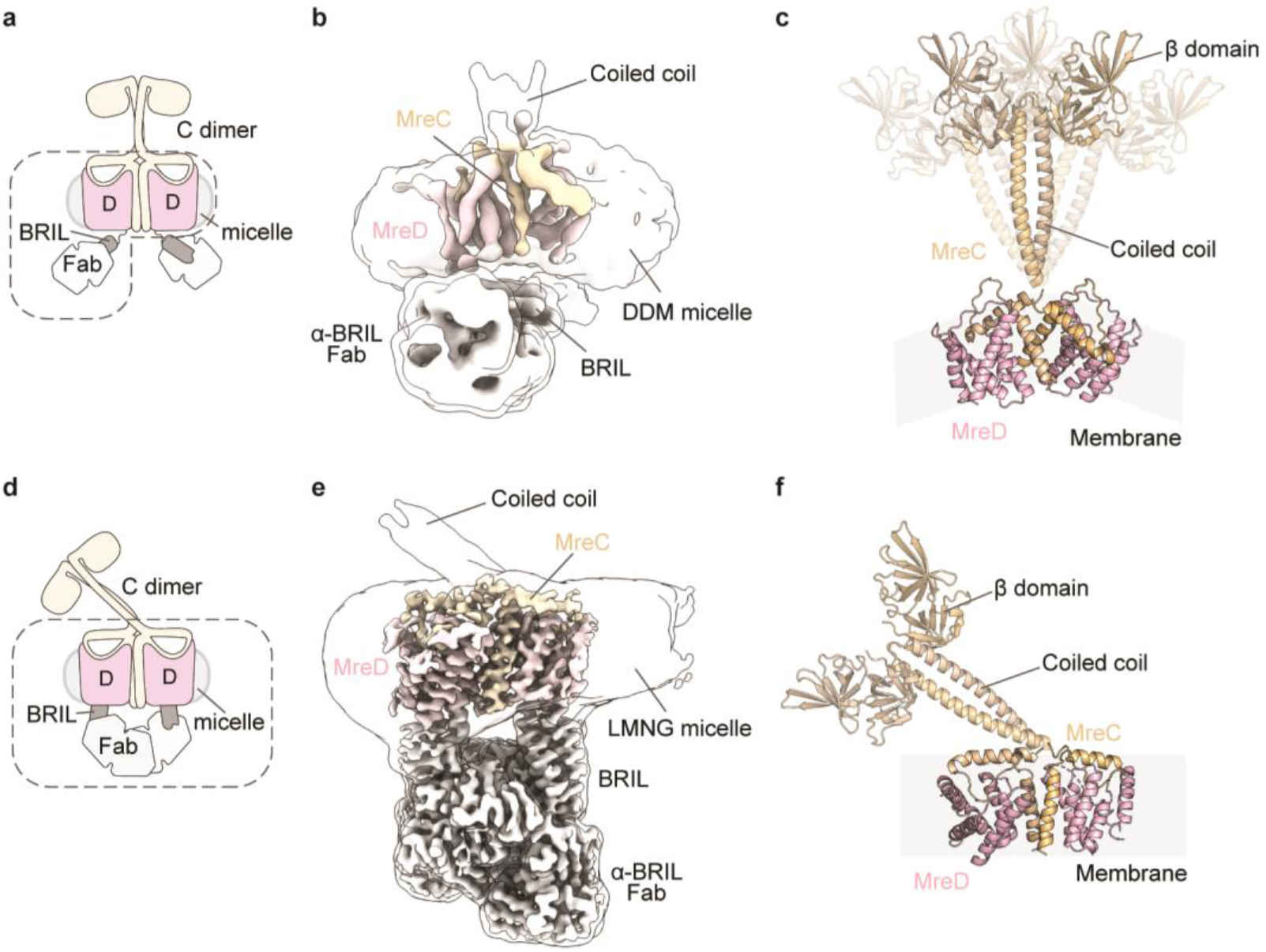
*Tt*MreC-MreD coiled coil domain adopts multiple tilted conformations. **a**, Schematic representation of the overall architecture of the TtMreC-MreD-BRIL (flexible C-terminal fusion) + α-BRIL Fab complex used in the low-resolution structure, colored as in Fig. 1a. The dashed line outlines the regions most clearly resolved in the cryo-EM map. **b**, The cryo-EM map of the *Tt*MreC and *Tt*MreD complex determined to 7.4 Å resolution is shown, colored according to the cartoon in (**a**). The same map, shown at a high threshold level and smoothed with gaussian filtering is overlaid in transparent surfaces to illustrate the relative position of the coiled coil domains of *Tt*MreC. **c**, Two copies of the transmembrane (TM) and membrane proximal regions of *Tt*MreC (wheat) and *Tt*MreD (pink) that were docked into the low-resolution map are shown in ribbons. The approximate position of the membrane is annotated in grey. Three copies of the two β domains and coiled coil of *Tt*MreC are shown as tan ribbons depicting possible orientations of this region based on the density observed in the cryo-EM map. **d-f**, The same panels as in a-c but for the 3.6 Å resolution structure of *Tt*MreC-MreD with the rigid internal fusion of BRIL. In **f**, only one copy of the β domains and coiled coil of *Tt*MreC is shown as tan ribbons, depicting the single tilted orientation observed in the map.

### MreC membrane proximal helix and cavity loop are required for Rod complex activity *in vivo*

Our *in vitro* data suggest that MreD-mediated changes to MreC conformation are required for activation of RodA-PBP2. To test this prediction, we employed a library screening approach to determine how perturbations to the membrane-proximal regions of MreC affect Rod complex function in *E. coli*. We designed a site saturation variant library (SSVL) in which all possible amino acid substitutions were incorporated at each position of the predicted cavity loop and membrane proximal helix of *Ec*MreC (residues 29-76; Supplemental Fig. 9a). We screened the resulting library using a previously described assay, in which loss of growth can be rescued by mutations that either directly or indirectly disrupt normal glycan strand synthesis by the Rod complex^25^. We identified 12 clones with disrupted Rod complex activity, of which 8 were selected for further validation (*Methods*, Supplemental Table 3).

We tested the activity of these mutants in dominant-negative and complementation assays. Seven of the 8 mutants had growth defects in one of the two assays. In the complementation assay, variants are tested for their ability to restore growth of the Δ*mreC* mutant. In the dominant negative assay, MreC mutants that retain the ability to bind MreD but fail to activate PG synthesis prevent cellular growth. Two cavity loop mutants (I38D and T44D) exhibited strong dominant-negative activity at moderate induction levels and failed to complement the growth defect of the mreC deletion mutant (Fig. 5). A different substitution (I38N) at the same position had attenuated dominant-negative activity but was still not functional in the complementation assay (Fig. 5). Although membrane-proximal helix mutants (V63Q, L59T, Y41K, M42F) did not show dominant-negative activity, all of them exhibited reduced growth in the complementation assay (Fig. 5). The V63Q substitution completely abolished growth, whereas L59T, Y41K, M42F variants had a mild growth defect. Importantly, these phenotypes were not due to reduced levels of MreC expression (Supplemental Fig. 9b,c). Collectively, these results demonstrate that the cavity loop and membrane-proximal helix of *Ec*MreC are both critical for Rod complex function *in vivo*, supporting a model in which MreD controls the conformation of MreC to regulate the activity of the Rod complex.

**Fig. 5.**
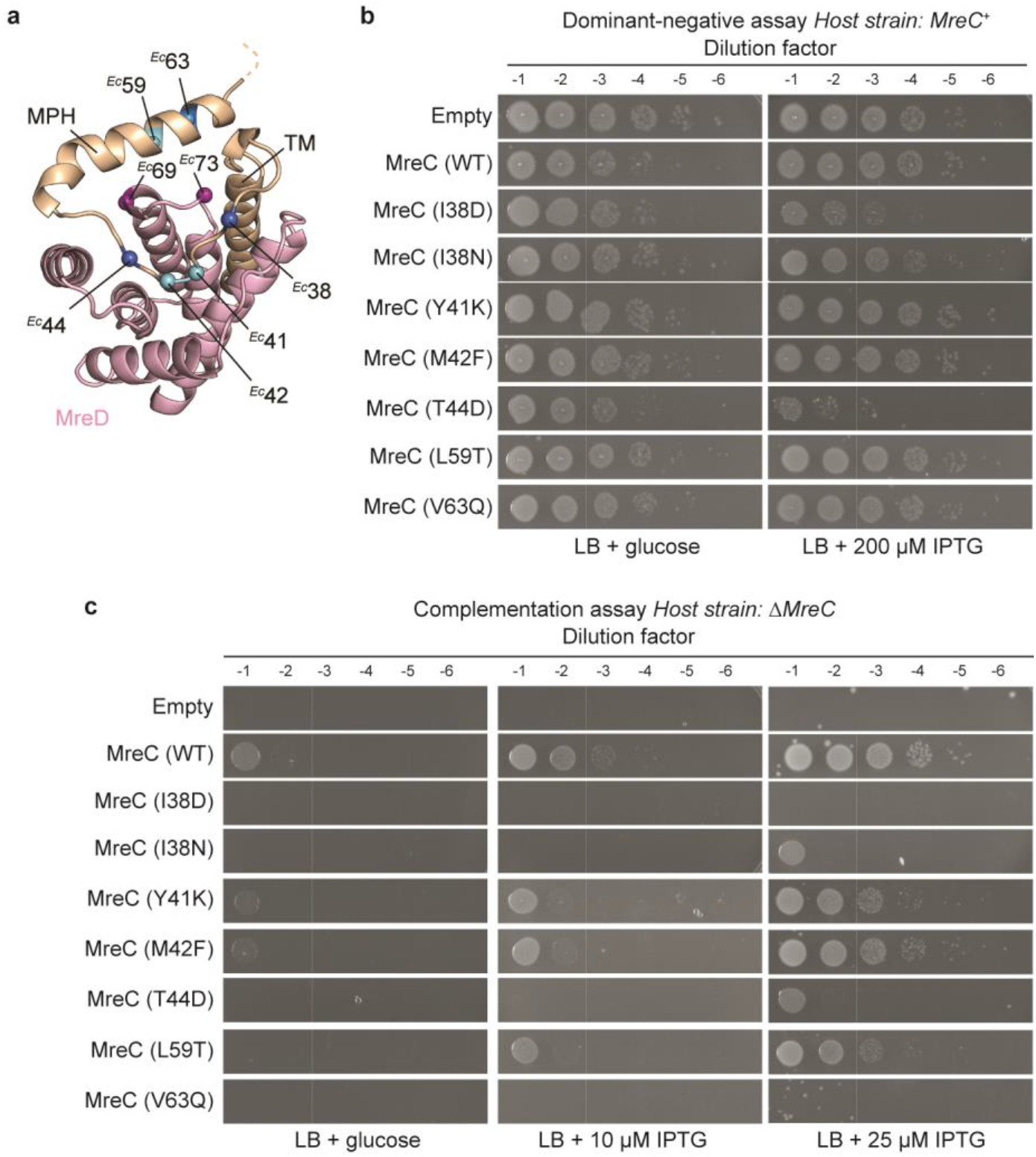
*Ec*MreC cavity loop and membrane proximal helix interfaces are critical for Rod complex activity *in vivo*. **a**, *Tt*MreC and *Tt*MreD are shown as cartoons colored tan and pink, respectively. The C-alpha carbons for the predicted positions of the *Ec*MreC mutants identified in the library screen are shown as spheres. Dark blue spheres indicate mutants with strong dominant-negative and/or complementation phenotypes and light blue spheres indicate a modest complementation defect. Purple spheres indicate previously identified dominant-negative alleles of *Ec*MreD. **b**, Cultures of wild-type *E. coli* (MG1655) transformed with Ptac vectors encoding each allele of *mreC* identified in the library screen were serially diluted and spotted on LB agar with either 0.2% glucose or 200 μM IPTG. **c**, Cultures of the Δ*mreC* strain of *E. coli* (MT4) transformed with P_lac_ vectors encoding the same alleles of *mreC* as in (**b**) were serially diluted and spotted on LB agar with either 0.2% glucose, 10 μM IPTG, or 25 μM IPTG.

## Discussion

The Rod complex has been extensively studied by genetic and cell imaging approaches, yet the mechanisms of its activity and regulation have remained unclear. Here, we resolve the first structure of the essential MreC-MreD complex and provide a model for its role in the activation of PG synthesis by the Rod complex (Fig. 6). We propose that in the apo state, MreC exists in an extended conformation that positions its β domain far above the bacterial membrane. Upon binding to MreD, contacts at the cavity loop and MPH interfaces introduce a break in the alpha helical domain of MreC that lowers the β domain and allows the coiled coil to undergo a rapid tilting motion. Once the MreC-MreD complex samples a state compatible with PBP2 binding, it is captured in this conformation by interactions between PBP2 and the β domain (Fig. 6). This predicted binding mode results in an assembly in which the two subcomplexes (RodA-PBP2 and MreC-MreD) do not interact within their transmembrane domains but are instead tethered by a single interface in the periplasm (Supplemental Fig. 8).

**Fig. 6.**
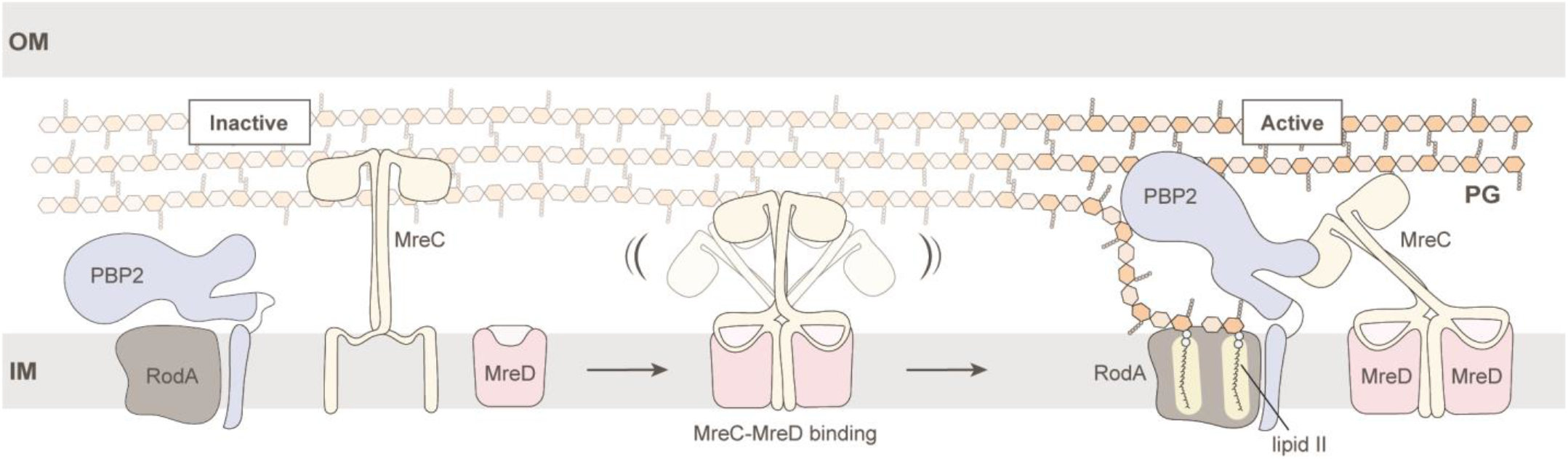
MreD modulates MreC conformation to facilitate Rod complex activation. Schematic overview shows the model for Rod complex activation. In the apo state of MreC, the β domains are elevated well above PBP2 and the bacterial membrane. The multiple MreC-MreD interfaces result in conformational changes in MreC that lower the β domains and activate tilting of the coiled coil region. In the tilted conformation, the β domain can now access its binding site on PBP2. This interaction results in allosteric opening of PBP2 and thereby activation of RodA.

Although the SEDS-bPBP pair of the cell division machinery (FtsWI) is structurally related to RodA-PBP2, the sequence, structure, and binding modes of their activators (FtsQLB) are highly divergent. The recently determined structure of the divisome (*Pa*FtsWI-FtsQLB) shows a transmembrane interaction between FtsWI and the heterodimeric FtsB-FtsL activators^13^. In addition, FtsB and FtsL form a coiled coil that directly interacts with FtsI, whereas MreC interacts with PBP2 via the β domain and seemingly relies on the coiled coil only for its proper positioning. Despite binding in different modes, the divisome regulators may trigger the same allosteric changes in the PG synthase—specifically, structural opening of FtsI—to promote activation. Further studies are needed to determine how the allosteric signals from regulatory protein binding are propagated to the active sites of the PG synthase.

Our cryo-EM structure demonstrates that MreD adopts an S-component fold, with a periplasmic-facing pocket analogous to the ligand binding site of these transporters^29^. In S components, micronutrients bind within this pocket, leading to protein toppling within the bacterial membrane and ligand transport into the cytoplasm^35-39^. In our structures, the two MreD molecules are tilted relative to the predicted membrane plane (Supplemental Fig. 10), suggesting that MreD may also undergo some degree of membrane toppling. Further studies will be required to determine what orientation MreD adopts in the native membrane and how such a toppling motion might influence its interaction with MreC.

Our work reveals a novel function for the S-component fold, in which the ligand binding site has been repurposed to recognize a protein ligand and serve as a conformational activation switch. We speculate that the MreD pocket may also recognize an as-yet unidentified small molecule ligand, which would compete with MreC for binding to MreD and have an inhibitory effect on Rod complex activity. The nature of this molecule is unclear, but attractive candidates are PG precursors or degradation products that might coordinate Rod complex activity with other aspects of cell envelope biogenesis. Regardless of whether this mode of regulation exists naturally, targeting the MreD pocket with a small molecule inhibitor offers a promising avenue for novel antibiotic development.

## Acknowledgements

Cryo-EM data were collected at the Harvard Cryo-EM Center for Structural Biology at Harvard Medical School. Structural biology applications used in this project were compiled and configured by SBGrid. M.S.A.G. is supported by the HHMI Hanna H. Gray Fellows Program. I.S. is supported by the Van Maanen Fellowship from the Department of Biological Chemistry and Molecular Pharmacology at Harvard Medical School. E.M.F. and I.S. are supported by the National Science Foundation (NSF) Graduate Research Fellowship award. Funding for this work was provided by National Institutes of Health grants U19 AI158028 (A.C.K. and T.G.B.), R01 GM114065 (J.J.L.), R01 AI083365 (T.G.B.), and R00GM135519 (N.F.P.), and investigator funds from the Howard Hughes Medical Institute (T.G.B.). N.F.P. acknowledges support from the Innovation Research Fund of the Dana-Farber Cancer Institute. We thank members of the Bernhardt, Loparo, and Kruse labs for their helpful suggestions and members of the Harvard Cryo-EM Center for Structural Biology and SBGrid for their support. This paper was typeset with the bioRxiv word template by @Chrelli: www.github.com/chrelli/bi-oRxiv-word-template

## Data Availability

The atomic coordinates for the *Tt*MreC-MreD-BRIL internal fusion + α-BRIL Fab and *Tt*MreC-MreD-BRIL flexible fusion + α-BRIL Fab structures have been deposited at the Protein Data Bank with accession codes 9DVB and 9DVC, respectively. Corresponding cryo-EM maps are deposited at the Electron Microscopy Data Bank and assigned codes EMD-47199 and EMD-47200, respectively. Coordinates and maps are available prior to PDB release upon request. Custom code is available upon request.

## Materials and Methods

### Production of anti-BRIL antibody fragment

The previously described anti-BRIL Fab (BAG2) heavy chain was cloned into pTarget^31^. We incorporated a set of previously described mutations in the hinge region of the Fab heavy chain (SSASTKG replaced with FNQIKG) to increase its rigidity and assist with particle alignment in cryo-EM^32^. The BAG2 light chain was cloned into pD261040. Heavy and light chain DNA was combined at a ratio of 1:1 (0.75 mg/l culture) and mixed with FectoPro (800 μL/l culture) in 100 ml OPTI-MEM for 10 min. The resulting mixture was used to transfect 0.9 l of Expi293F TetR cells at a density of ∼3.2 × 10^6^ cells/ml. Approximately 24 hrs later, 6 mM valproic acid and 0.8% glucose were added to the cultures. After 7 days, the cellular supernatant was harvested and 200 ml 1M HEPES pH 8.0 and 100 ml 5 M NaCl were added. Fab was purified from this solution using CaptureSelect CH1-XL Affinity Matrix (ThermoScientific) resin which had been equilibrated in HEPES buffered saline (HBS). The resin was washed twice using 2 CVs of HBS and Fab was eluted using 0.1 M glycine pH 3.0 which was neutralized using a 20 X stock solution of HBS. The final Fab was exchanged into HBS by either dialysis or size-exclusion chromatography (SEC).

### Design of MreD-BRIL Fusion

We created the MreD-BRIL fusion using a structural bioinformatics approach^41^. The method searches the PDB to extend two helices of MreD (AlphaFold prediction), onto which BRIL is superimposed. We then designed the sequence of the linking helical regions using proteinMPNN^42^.

### Protein expression and purification for *Tt*MreC-MreD complexes

Plasmids encoding *T. thermophilus* MreC and variants thereof (pMG-7, pMG-113, pMG-114, pMG-126, pMG-127, pMG-128) or MreD (pDSG-7, pMG-54, pMG-117, pMG-116) (Supplemental Table 4) were transformed into *E. coli* C43 (DE3) cells with or without SUMO tag-specific Ulp1 protease under an arabinose inducible plasmid (pAM174) respectively. Following transformation, bacteria were grown on Lennox broth (LB) agar plates with 100 µg/ml ampicillin (amp), 35 µg/ml chloramphenicol (cam), and 50 µg/ml spectinomycin (spec) at 37 °C for ∼16 h. Transformants were inoculated into 5 ml Terrific broth (TB) + amp, spec, cam and grown for 16 h at 37 °C with shaking. These cultures were diluted into 1 l of TB + amp, spec, cam, 2 mM MgCl_2_, 0.1% glucose, and 0.4% glycerol, and grown at 37 °C with shaking until an OD600 >2. Cells were then cooled to 18 °C while shaking and, after 1 h, protein expression was induced by adding 1 mM isopropyl β-d-1-thiogalactopyranoside (IPTG) and/or 0.1% arabinose to the culture. After ∼16 h of induction at 18 °C, bacteria were harvested by centrifugation at 5,000 x g for 30 min and the pellets were flash frozen and stored at -80 °C for subsequent purification.

Frozen bacterial pellets were resuspended by stirring in lysis buffer (50 mM HEPES pH 7.5, 150 mM NaCl, 20 mM MgCl_2_) with added protease inhibitors tablets (ThermoFisher) and benzonase nuclease (∼1.5 units/ml, Sigma Aldrich) at a ratio of 5 ml of buffer per gram of pellet. Resuspended bacteria were passed through a Dounce homogenizer and afterwards lysed by four passes through an LM10 microfluidizer (Microfluidics) at 17,000 psi. The bacterial lysate was then centrifuged for 1 h at 50,000 x g to separate membrane (pellet) and cytoplasmic (supernatant) cellular fractions. After discarding the supernatant, solubilization buffer (20 mM HEPES pH 7.5, 0.5 M NaCl, 5% glycerol, 1% n-dodecyl-β-D-maltopyranoside (DDM)) with protease inhibitor tablets was added to the membrane fraction (150 ml buffer/1 l culture) and the fraction was homogenized using an UltraTurrax T25 electronic homogenizer (IKA) for 20 s at 13,000 rpm. This membrane mixture was stirred for ∼2 h at 4°C and centrifuged at 50,000 x g for 1 h to separate soluble (supernatant) and insoluble fractions. The supernatant was passed through a fiberglass filter to remove large particles and supplemented with 2 mM CaCl_2_ prior to loading onto affinity resin.

In the case of *Tt*MreC-MreD complexes, the filtered supernatant supplemented with 2 mM CaCl_2_ was first loaded onto an anti-FLAG M1 agarose resin column previously equilibrated into solubilization buffer supplemented with 2 mM CaCl_2_. After supernatant application, the column was washed with 2 CVs of solubilization buffer with 2 mM CaCl_2_, followed by 3 washes with 2 CVs of wash buffer (20 mM HEPES pH 7.5, 500 mM NaCl, 5% glycerol, 2 mM CaCl2, 0.1% DDM). Protein was eluted with ∼5 CVs of FLAG elution buffer (20 mM HEPES pH 7.5, 500 mM NaCl, 5% glycerol, 0.1% DDM, 0.4 mg/ml FLAG peptide). This elution was supplemented with 2 mM CaCl_2_ and then loaded onto an anti-Protein C tag agarose resin column pre-equilibrated with wash buffer. After passing the FLAG elution 3 times over the anti-Protein C column, the column was washed 3 times with 2 CVs of wash buffer (20 mM HEPES pH 7.5, 500 mM NaCl, 5% glycerol, 2 mM CaCl_2_, 0.1% DDM) and eluted using ∼ 5 CVs of Protein C elution buffer (20 mM HEPES pH 7.5, 500 mM NaCl, 5% glycerol, 0.1% DDM, 0.2 mg/ml Protein C peptide, 5 mM EDTA pH 8). For cryo-EM samples containing the rigidly fused BRIL (pDSG-7), the sample was equilibrated stepwise into lauryl maltose neopentyl glycol (LMNG) while immobilized on the anti-Protein C column using 2 washes each of 3 CVs of wash buffer with 0.1% DDM/0.1% LMNG, 0.01% DDM/0.1% LMNG, and 0.1% LMNG. The protein was eluted using 5 CVs of Protein C elution buffer (20 mM HEPES pH 7.5, 500 mM NaCl, 5% glycerol, 0.1% LMNG, 0.2 mg/ml Protein C peptide, 5 mM EDTA pH 8). Eluates were concentrated to ∼ 500 µl using an Amicon Ultra Centrifugal filter (Millipore Sigma) with a 50 kDa molecular weight cutoff (MWCO) for samples lacking the anti-BRIL Fab and a 100 kDa MWCO for samples in which the Fab was present. For samples containing the anti-BRIL Fab, at least 1.5-fold molar excess of the Fab was added to the sample and incubated for ∼10 min at room temperature to allow complex formation. The samples were additionally purified using an Superdex 200 Increase (Cytiva) SEC column. For cryo-EM samples containing the rigidly fused BRIL (pDSG-7), LMNG SEC buffer was used (20 mM HEPES pH 7.5, 350 mM NaCl, 0.01% LMNG). For cryo-EM samples containing the flexibly fused BRIL MreD construct (pMG-54), low DDM SEC buffer was used (20 mM HEPES pH 7.5, 350 mM NaCl, 0.02% DDM). For samples used in truncation analysis (pMG-113, pMG-114, pMG-126, pMG-127, pMG-128, pMG-54, pMG-117, pMG-116), DDM SEC buffer was used (20 mM HEPES pH 7.5, 350 mM NaCl, 0.1% DDM).

### Construct design for smFRET microscopy

All constructs used for smFRET microscopy were assembled via PCR and Gibson assembly (Supplemental Table 4). To ensure that only one residue pair was fluorescently labeled within the *Tt*MreC-MreD complex, *Tt*MreC was monomerized by replacing part of its coiled coil (*Tt*MreC^70-96^) with a monomeric stable alpha helical domain (SAH) based on a murine myosin X peptide, as previously discussed^43^. Apo MreC constructs contained an N-terminal ALFA-tag for immobilization on streptavidin-functionalized coverslips via a biotinylated α-ALFA nanobody (NanoTag). To enable tandem purification of stoichiometric *Tt*MreC-MreD complex, a Protein C tag was added to the *Tt*MreC C-terminus while an N-terminal SUMO-FLAG tag system was added to *Tt*MreD. The *Tt*MreC-MreD complex was similarly captured in the flow cell via an N-terminal ALFA tag preceding the FLAG tag in MreD.

### smFRET sample preparation and labeling

*Tt*MreC (pSI214, pSI291, pSI292, pSI324, pSI325, pSI329, pDSG18) and *Tt*MreD (pMG54, pDSG22, pDSG26) smFRET constructs were expressed and purified as above until the affinity purification step, where all the samples were washed and eluted in 0.1% DDM. These respective eluates were concentrated to ∼ 500 µl using an Amicon Ultra Centrifugal filter (Millipore Sigma) with a 50 kDa molecular weight cutoff (MWCO). To prevent oxidation, concentrated protein was reduced with 10 mM DTT for 15 min at room temperature and then loaded onto a Superdex 200 Increase (Cytiva) size exclusion chromatography (SEC) column in SEC buffer (20 mM HEPES pH 7.5, 350 mM NaCl, 0.1% DDM, 5 mM EDTA). Fractions corresponding to monomeric apo *Tt*MreC or the *Tt*MreC-MreD complex were collected and concentrated using a 50 kDa MWCO filter for fluorescent labeling. Samples were labeled by incubation with 10-fold molar excess of sulfo-Cyanine3-maleimide (Cy3, Lumiprobe) and sulfo-Cyanine5-maleimide (Cy5, Lumiprobe) for 30 min at room temperature in SEC buffer. Unreacted fluorophores were removed by passing labeled protein through a 2 ml Zeba spin desalting column (Thermo Fisher Scientific). Following Zeba exchange, the labeled proteins were purified by SEC as described above and monomeric fractions were collected, concentrated, flash frozen, and stored at -80 °C for subsequent smFRET imaging.

### smFRET chamber preparation and data collection

Microfluidic chambers for smFRET imaging were built as previously described^24,44,45^. Briefly, a glass coverslip was functionalized with methoxypolyethylene glycol-succinimidyl valerate MW 5000 (mPEG-SVA-5000, Laysan Bio Inc.) and biotin-methoxypolyethylene glycol-succinimidyl valerate MW 5000 (biotin-PEG-SVA-5000) and stored for up to two weeks at -20°C before use. Strips of double-sided tape were then sandwiched between a quartz slide and the coverslip with polyethylene tubing attached (PE #20 and PE #60, VWR), and epoxy was used to seal these layers together to form a microfluidic chamber. Solutions were added to the chamber through a 1 ml syringe (VWR).

Prior to imaging, the chamber was blocked with 1 mg/ml bovine serum albumin solution (NEB) for 5 min, followed by two washes with SEC buffer. Next, the surface was functionalized with 0.25 mg/ml streptavidin for 5 min, followed by two washes with SEC Buffer to remove unbound streptavidin. Next, 0.1 mg/ml anti-ALFA tag biotinylated nanobody was added, incubated for 5 mins, and excess nanobody was washed off with SEC buffer. Following this, labeled protein was added to the chamber at 100-500 pM concentration and incubated with a reactive oxygen species scavenging cocktail consisting of SEC buffer with 5 mM protocatechuic acid (PCA), 0.1 µM protocatechuate-3,4-dioxygenase (PCD), 1 mM ascorbic acid (AA), and 1 mM methyl viologen (MV) for 5 min.

Images were collected on an Olympus IX-71 total internal reflection (TIRF) microscope using Hamamatsu HCImage live version 4.4.0.1 and Labview version 15.0f2 software as previously described^24,44^. Power was set to 2 W/cm2 for the 532 nm laser and 1 W/cm^2^ for the 641nm laser. For each sample, movies of 60-90 seconds in length were collected at a frame rate of 4 s^-1^ with two frames of 532 nm excitation alternating with one frame 641 nm excitation.

### smFRET analysis

Raw movies were parsed using custom MATLAB scripts as previously described to isolate single-molecule trajectories^24,44^. Importantly, we used co-localization between donor and acceptor foci, rather than their FRET signal, to identify doubly labeled complexes and ensure that trajectories with FRET ∼ 0 were not excluded. Trajectories were subsequently filtered to include only complexes with a single donor and a single acceptor and cropped at the time of photobleaching using a custom Matlab script. Filtered trajectories were then analyzed using the ebFRET GUI to quantify the kinetics and distributions of the underlying smFRET states^46^.

### Cryo-EM sample preparation and data collection

The *Tt*MreC-MreD-BRIL (internal) + anti-BRIL Fab sample in LMNG (pMG-7, pDSG-7, pMG-36, pMAS-266) was concentrated to 4.7 mg/ml. C-flat grids with 1.2-μm diameter/1.3-μm spacing (Electron Microscopy Sciences) were plasma cleaned for 30 sec. Grids were prepared using a Vitrobot Mark IV (ThermoFisher) using 4 ul sample with a 10 s wait time and a 5 s blot time at force 15 before plunging into liquid ethane. Data were collected on a Talos Arctica operating at 200 kV (ThermoFisher Scientific) equipped with a K3 direct electron detector (Gatan) using Serial EM at the Harvard Cryo-Electron Microscopy Center for Structural Biology. The *Tt*MreC-MreD-BRIL (C-terminal) + anti-BRIL Fab sample in DDM (pMG-7, pMG-54, pMG-36, pMAS-266) was prepared similarly, except that the sample was concentrated to 3.6 mg/ml and grids were blotted for 8 s. See Supplemental Table 1 for data collection parameters.

### Cryo-EM data processing and model building

Motion correction and dose weighting, followed by patch-based contrast transfer function (CTF) estimation, blob picking, and local motion correction were performed in CryoSPARC Live^47,48^. For the *Tt*MreC-MreD-BRIL (internal) + anti-BRIL Fab dataset, 2D classification on the initial particle stack was used to generate templates used in template-based picking using CryoSPARC. *An initio* models were also generated from the initial blob-picked particle stack. The template picked particles were subjected to multiple rounds of heterogeneous refinement in CryoSPARC. A stack of 206,722 particles was subjected to non-uniform refinement^49^, reference-based motion correction and global CTF refinement. Particle rebalancing resulted in a stack of 166,346 particles, which were refined to a global resolution of 3.6 Å. Local resolution estimation and filtering was carried out in CryoSPARC. Although masking to remove the detergent micelle improved the nominal resolution, the quality of the map was highest when using automatic mask generation in non-uniform refinement. The coordinates were built manually in Coot^50^, using starting models generated from a combination of AlphaFold2 predictions and the existing BRIL + anti-BRIL Fab structure^31,33^. The structure was refined using real-space refinement in Phenix^51,52^.

For the *Tt*MreC-MreD-BRIL (C-terminal) + anti-BRIL Fab sample, pre-processing was carried out as described above. Two rounds of 2D classification were carried out in CryoSPARC. The resulting stack of 206,339 particles was used for *ab initio* model generation. Multiple rounds of heterogeneous refinement resulted in a stack of 51,857 particles. A mask including a single copy of *Tt*MreC-MreD-BRIL and one anti-BRIL Fab was generated in Chimera^53^ and used for local refinement in CryoSPARC, resulting in a 7.4 Å resolution map. The orientation of the two heterodimers relative to one another differed between this map and the higher resolution map determined above (Supplemental Fig. 10). Therefore, two individual *Tt*MreC-MreD dimers were docked into this map separately, without additional model building. Structural biology software used in this project was compiled and configured by SBGrid^54^. Figures were generated using PyMOL and ChimeraX^55^.

### *In vivo* library screen

A previously described high-throughput screen was used to identify defective mutants of *Ec*MreC that were not defective due to changes in protein expression or Rod complex incorporation^25^. *Ec*MreC contains an inhibitory gamma domain and mutations in this region are associated with strong dominant-negative phenotypes^25^. To avoid selection of gamma domain mutants, we focused our screen only on the MreD-proximal regions of MreC. This assay exploits the fact that Rod complex activity is non-essential in *E. coli* when FtsZ is overexpressed^20^, allowing for survival under conditions in which the Rod system is defective. Mecillinam, an antibiotic that blocks crosslinking by PBP2, remains toxic in this background because it induces futile cycles of glycan strand synthesis and turnover^56^. In this context, mecillinam treatment selects for mutants of MreC that incorporate into the Rod complex and displace functional copies of MreC but fail to activate glycan strand synthesis by RodA.

A 960-member site saturation variant library (SSVL) library in which all possible amino acid substitutions were incorporated at each position of the predicted cavity loop and membrane-proximal helix of *Ec*MreC (residues 29-76) was synthesized as double-stranded DNA (Twist Bioscience). These DNA fragments were then cloned into the pPR11 background, in which the *mreCD* operon is expressed under the IPTG (isopropyl-β-d-thiogalactopyranoside)-inducible tac promoter (P_tac_). The fragments and the pPR11 plasmid were digested with BamHI and EcoRI (NEB) and ligated with T4 ligase (NEB) before being transformed into NEB DH5 alpha cells and plated on LB-Cam35. The resulting colonies were scraped and pooled, then used to inoculate a culture for plasmid isolation. Wild-type *E. coli* (MG1655) cells were transformed with pTB63 encoding ftsQAZ^20^. Competent cells (MG1655-pTB63) were generated using transformation and storage solution^57^ and used for library and control vector transformations. Transformations were plated in duplicate on LB-Cam35-Tet5 (Control) and LB-Cam^35^-Tet^5^ + 50 μM IPTG + 2.5 μg/ml mecillinam. Control plates were used to estimate the total number of transformants as >80,000, indicating that the library diversity had been sampled more than 80-fold.

More than 40 colonies were re-streaked in duplicate on plates containing either mecillinam or mecillinam with IPTG to assess induction-dependence of antibiotic resistance. Eleven clones with varying degrees of IPTG-dependent mecillinam resistance were selected for sequencing. Of these, 4 had acquired additional mutations in the gamma domain and were excluded from further analysis. One mutant contained a cysteine substitution (V46C) and was excluded to remove possible confounding effects of cysteine modification, and another contained two substitutions (M42F/V63Q), which were isolated for subsequent validation experiments. Two mutants with substitutions at position 38 were identified independently (I38D and I38N).

### Dominant-negative assay

The individual mutations identified in the screen above were cloned into the pPR11 background (P_tac_). Wild-type *E. coli* (MG1655) cells were transformed with each plasmid and plated on LB-Cam^35^ with 0.2% glucose (LB-Cam-Glu). Plasmids used were pHC800 [empty vector], pPR11 [mreC(WT)], pMSG7 [mreC(I38D)], pMSG8 [mreC(I38N)], pMSG9 [mreC(Y41K)], pMSG10 [mreC(M42F)], pMSG11 [mreC(T44D)], pMSG12 [mreC(E58Y)], pMSG13 [mreC(L59T)], and pMSG14 [mreC(V63Q)], see Supplemental Table 5. Single colonies for each were re-streaked on LB-Cam-Glu before being used to inoculate overnight cultures. The OD600 of each overnight culture was normalized to OD_600_ = 1 using LB-Cam-Glu before being diluted serially 10-fold using LB. For each dilution, 4 μl was plated on media containing either LB, LB + glucose, or LB + IPTG (100 μM, 200 μM, 300 μM, 500 μM, or 1mM). All incubations were carried out at 30 °C.

### Complementation assay

The individual mutations identified in the screen above were cloned into the pMS5 background, in which the *mreCD* operon is expressed under the control of the IPTG-inducible lac promoter (P_lac_). *E. coli* cells in which the *mreC* gene had been deleted (MT4 (MG1655 ΔlacIZYA::frt mreC::kan))^16^ were transformed with each plasmid and plated on M9-Cam35 with 0.2% glucose (M9-Cam-Glu). Plasmids used were pPR66 [empty vector], pMS5 [mreC(WT)], pMSG17 [mreC(I38D)], pMSG18 [mreC(I38N)], pMSG19 [mreC(Y41K)], pMSG20 [mreC(M42F)], pMSG21 [mreC(T44D)], pMSG22 [mreC(E58Y)], pMSG23 [mreC(L59T)], and pMSG24 [mreC(V63Q)], see Supplemental Table 5. Single colonies for each were re-streaked on M9-Cam-Glu before being used to inoculate liquid cultures. The OD_600_ of each culture was normalized to OD_600_ = 1 using M9-Cam-Glu before being diluted serially 10-fold using M9. For each dilution, 4 μl was plated on media containing either LB, LB + glucose, or LB + IPTG (10 μM, 25 μM, 50 μM, 100 μM, 200 μM or 1 mM). All incubations were carried out at 30 °C.

### Western blot

Plasmids used for complementation assay were transformed into MT4 cells and plated on (M9-Cam-Glu). Single colonies for each were restreaked before being used to inoculate liquid cultures of M9-Cam35 with 0.2% maltose. After ∼48 hrs, these cultures were diluted 1:200 into M9-Cam^35^ with 0.2% maltose containing either 10 μM or 25 μM of IPTG and cultured until OD_600_ = 0.3-1.2. The equivalent of 5 OD_600_ units for each culture was centrifuged at 2000 x g for 10 min and pellets were frozen at - 20 °C for at least 20 min. Each pellet was resuspended in a solution composed of 50 mM HEPES pH 7.5, 150 mM NaCl, 2 mM MgCl_2_ and 0.25 units/μl benzonase and held at room temperature for at least 3 min. This sample was mixed in equal volume with 2X SDS loading buffer (6.32% SDS, 158 mM Tris-Cl, pH 6.8, 26.3% glycerol and 0.021% bromophenol blue) and incubated at room temperature for at least 30 min. Samples were separated using SDS-PAGE and transferred to a PVDF low fluorescence membrane using the Trans-Blot Turbo Transfer System (Bio-Rad). The membrane was blocked using 2% non-fat milk in tris-buffered saline with 0.1% Tween 20 (TSBT) for 1 hr at room temperature, rinsed with TBST, then incubated with primary antibody solution (anti-MreC antibody^25^ diluted 1:5,000 in 0.2% non-fat milk in TBST) for 1 hr at room temperature. The membrane was then washed 3 times with TBST, incubated with secondary antibody solution (IRDye 800CW Anti-Rabbit IgG Goat Secondary Antibody, LI-COR Biosciences) for 30 min at room temperature and washed again 3 times with TBST. The membrane was visualized using a fluorescence imager (Amersham Typhoon 5).

### AlphaFold3 modelling of four-component complex

Predictions of the Rod complex using AF3 (*Tt*MreC-MreD-RodA-PBP2) predict interactions that are free from the planar constraints of the membrane and did not accurately predict the MreC-MreD interface observed in the cryo-EM structure determined here. However, the predicted interaction between *Tt*RodA, *Tt*PBP2, and the β domain of *Tt*MreC agrees well with existing structural data for these subcomplexes^17,24,28^. We therefore combined these components of the AF3 model with our experimental data to generate a model for the four-component complex. First, we isolated the *Tt*MreC coiled coil domain with a single copy of the β domain from the AF3 structure (model 1). We manually positioned this subcomplex into the experimentally determined cryo-EM map to determine their approximate position relative to the experimentally determined transmembrane domains of TtMreC-MreD (model 2). We removed *Tt*MreD and all domains except for the PBP2-bound β domain of *Tt*MreC from the AF3 model. The remaining *Tt*RodA-PBP2-MreC β domain model (model 3) was combined with model 2, assuming a flexible linkage between the coiled coil and β domain of *Tt*MreC, while imposing an approximately planar orientation of the transmembrane domains of all four components. The resulting model did not predict any interaction between the transmembrane domains of the two subcomplexes (*Tt*RodA-PBP2 and *Tt*MreC-MreD).

**Supplemental Fig. 1.**
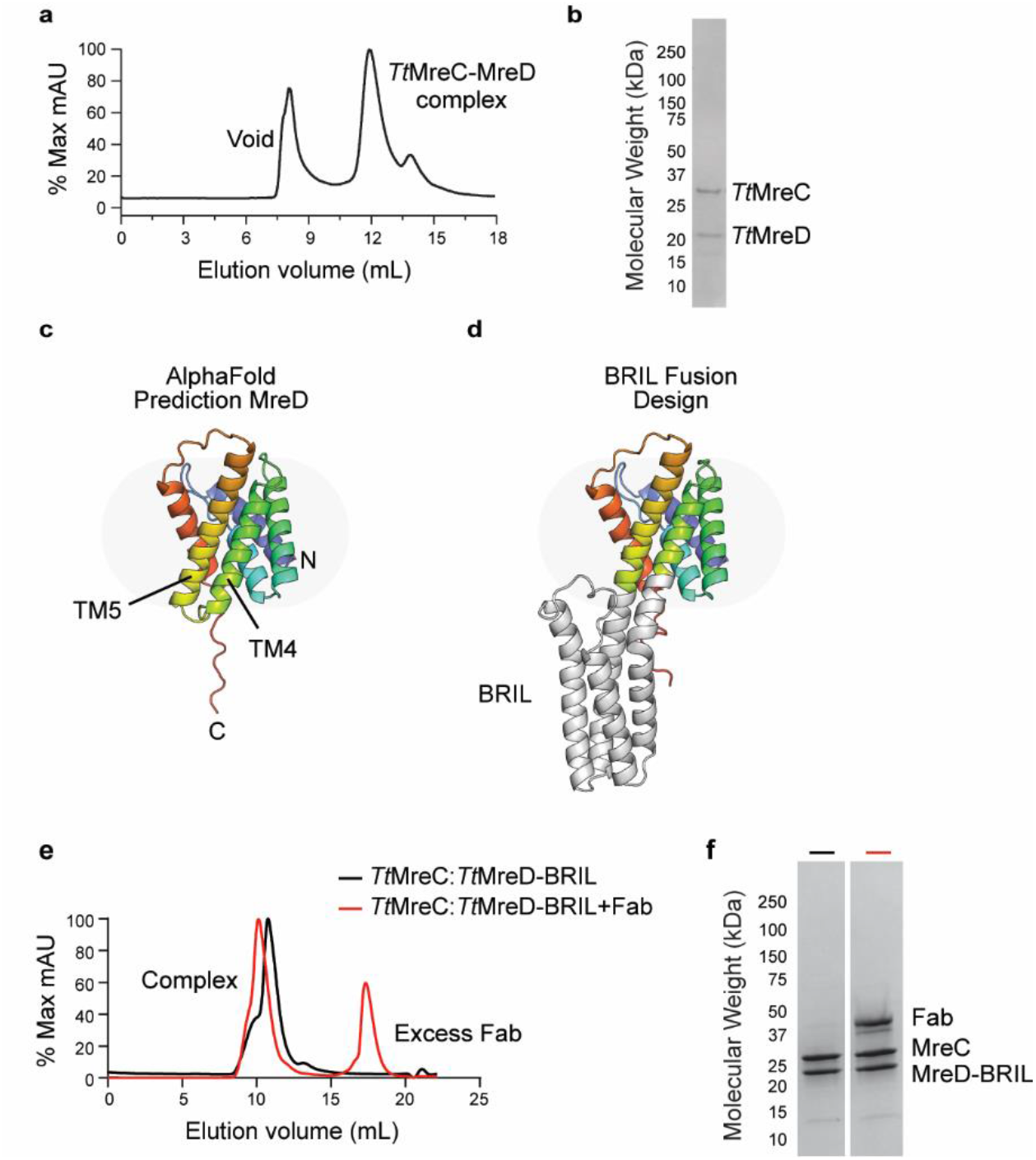
*Tt*MreC-MreD form a stable complex that is not disrupted by BRIL fusion. **a**, Size-exclusion chromatography (SEC) trace for the *Tt*MreC-MreD complex extracted with DDM and purified using sequential anti-M1 Flag and anti-Protein C affinity resins. **b**, SDS-PAGE of the resulting complex demonstrates that both proteins are present in the primary peak fractions on SEC. **c**, The AlphaFold structural prediction of *Tt*MreD is shown as ribbons colored in rainbow from the N-to C-terminus. **d**, The AlphaFold structural prediction of the final *Tt*MreD-BRIL is shown as ribbons colored as in (**c**). **e**, SEC trace and **f**, SDS-PAGE for the *Tt*MreC-MreD-BRIL complex with or without the α-BRIL Fab.

**Supplemental Fig. 2.**
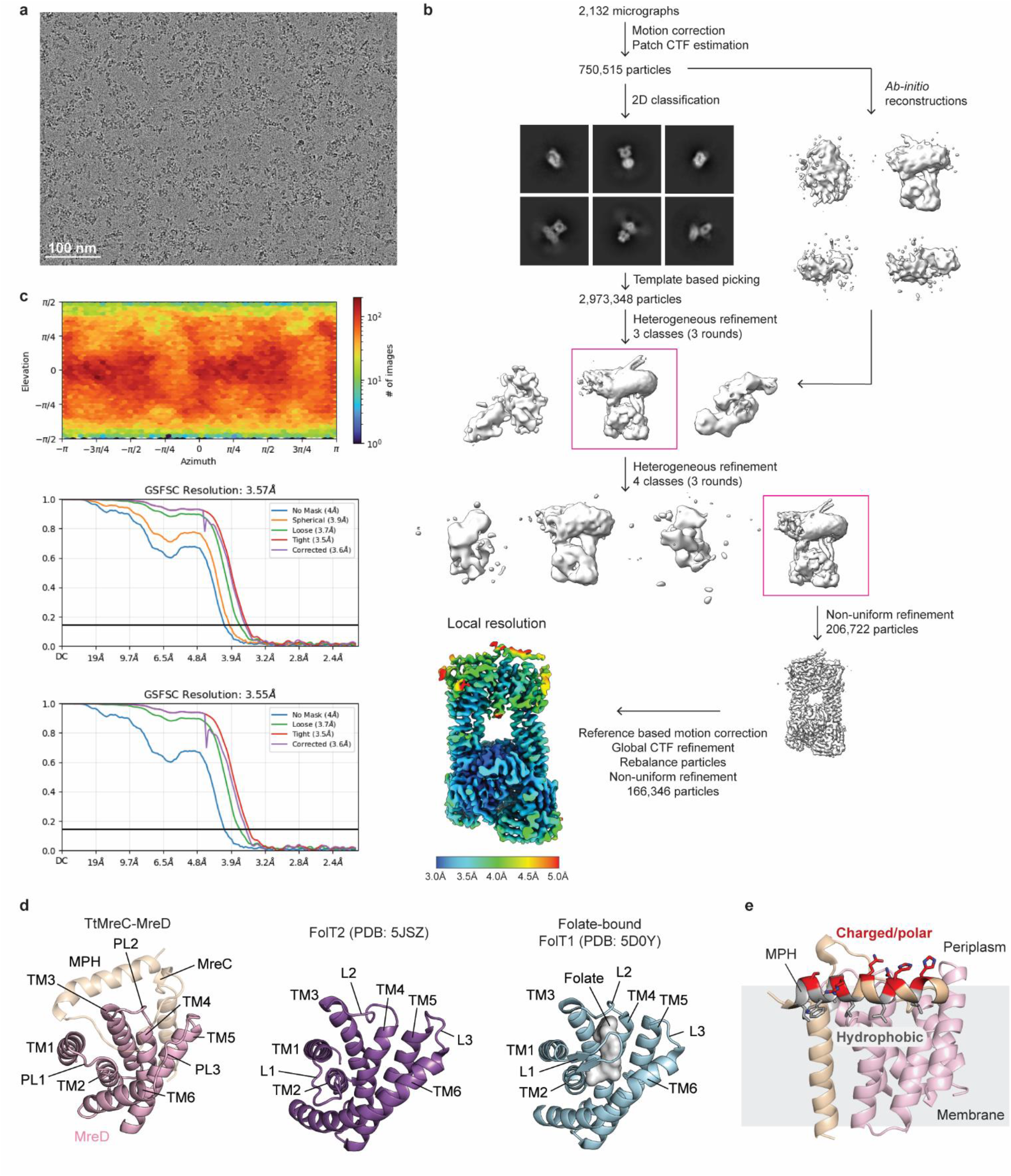
Cryo-EM data processing and analysis for the *Tt*MreC-MreD structure. **a**, Cryo-EM micrograph of the *Tt*MreC-MreD-BRIL + α-BRIL Fab in LMNG collected on a Talos Artica microscope. **b**, Processing scheme used in CryoSPARC to determine the complex structure and the resulting local resolution estimates for the final map. **c**, Viewing direction distribution plot (top), Gold-standard Fourier shell correlation (FSC) curves calculated in CryoSPARC before (middle) and after (bottom) FSC mask auto-tightening. The resolution was determined at FSC = 0.143 (horizontal black line). The final corrected mask gave an overall resolution of 3.6 Å. **d**, The structure of *Tt*MreC-MreD is shown as cartoons colored tan and pink. TM and PL refer to the transmembrane helices and periplasmic loops, respectively. The structure of the FolT2 S component was determined in the apo conformation, adopted when FolT2 is in complex with the ECF module of the folate transporter. FolT2 is shown as purple cartoons with the TMs and PLs labeled. The structure of the folate-bound FolT1 S component is shown as light blue ribbons with folate depicted as white molecular surfaces. TM6 is longer in the FolT structures than in *Tt*MreD and TM1 is ∼4 Å closer to the other TMs in FolT than in *Tt*MreD. In the ligand bound state, the first soluble loop of FolT1 rearranges to cap the folate binding site. The conformation of this region in *Tt*MreD more closely resembles the apo FolT2 structure. **e**, *Tt*MreC-MreD is shown as cartoons colored tan and pink, respectively, viewed rotated 180° relative to Fig. 1b. The membrane-proximal helix (MPH) residues with charged or polar characteristics are shown as red sticks. Hydrophobic MPH residues are shown as grey sticks. Oxygen and nitrogen atoms are colored red and blue, respectively. The approximate position of the membrane is annotated in grey.

**Supplemental Fig. 3.**
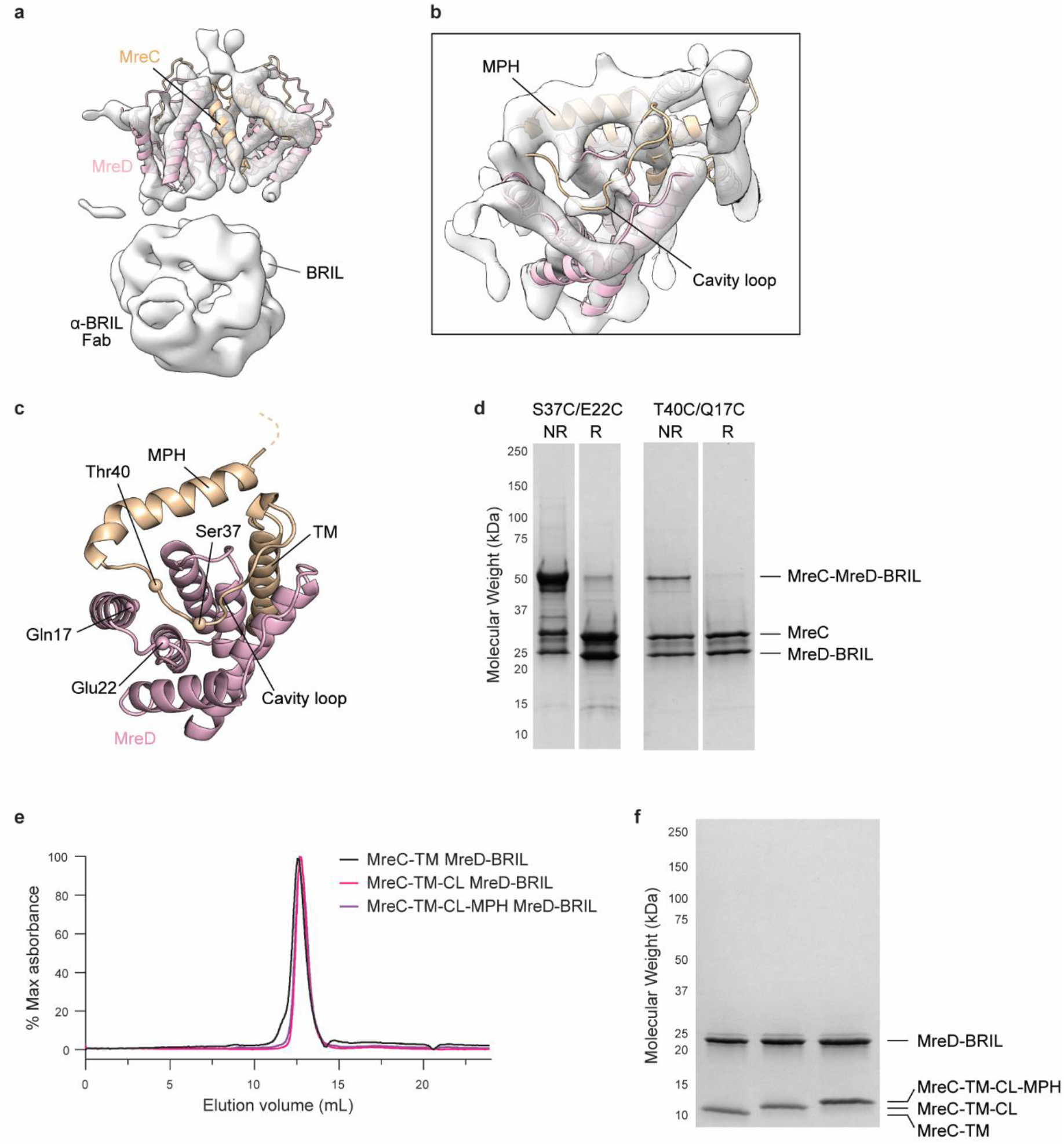
Structural and biochemical validation of the cavity loop and MPH interactions of *Tt*MreC-MreD. **a**, The 7.4 Å resolution cryo-EM map of the *Tt*MreC-MreD-BRIL (flexible C-terminal fusion) + α-BRIL Fab is shown in white. Two copies of the *Tt*MreC-MreD complex were docked into the density. **b**, Top view of the map shown in (**a**). The MPH of *Tt*MreC is clearly visible in the map. Weak density is also visible for the cavity loop. **c**, *Tt*MreC-MreD is shown as cartoons colored tan and pink, respectively. The C-alpha carbons of the residues mutated to cysteine are shown as spheres. **d**, SDS-PAGE shows the presence of disulfide crosslinked *Tt*MreC-MreD at ∼50 kDa. The presence of this band was dramatically reduced when the samples were incubated with β-mercaptoethanol to reduce disulfide bonds prior to separation. **e**, Three variants of *Tt*MreC were co-purified with *Tt*MreD-BRIL and formed a stable complex as measured by SEC and **f**, SDS-PAGE. MreC-TM included only the transmembrane (TM) domain (residues 1-28). MreC-TM-CL included the TM domain and the cavity loop (residues 1-45). MreC-TM-CL-MPH was composed of the TM domain, the cavity loop, and the membrane-proximal helix (residues 1-63).

**Supplemental Fig. 4.**
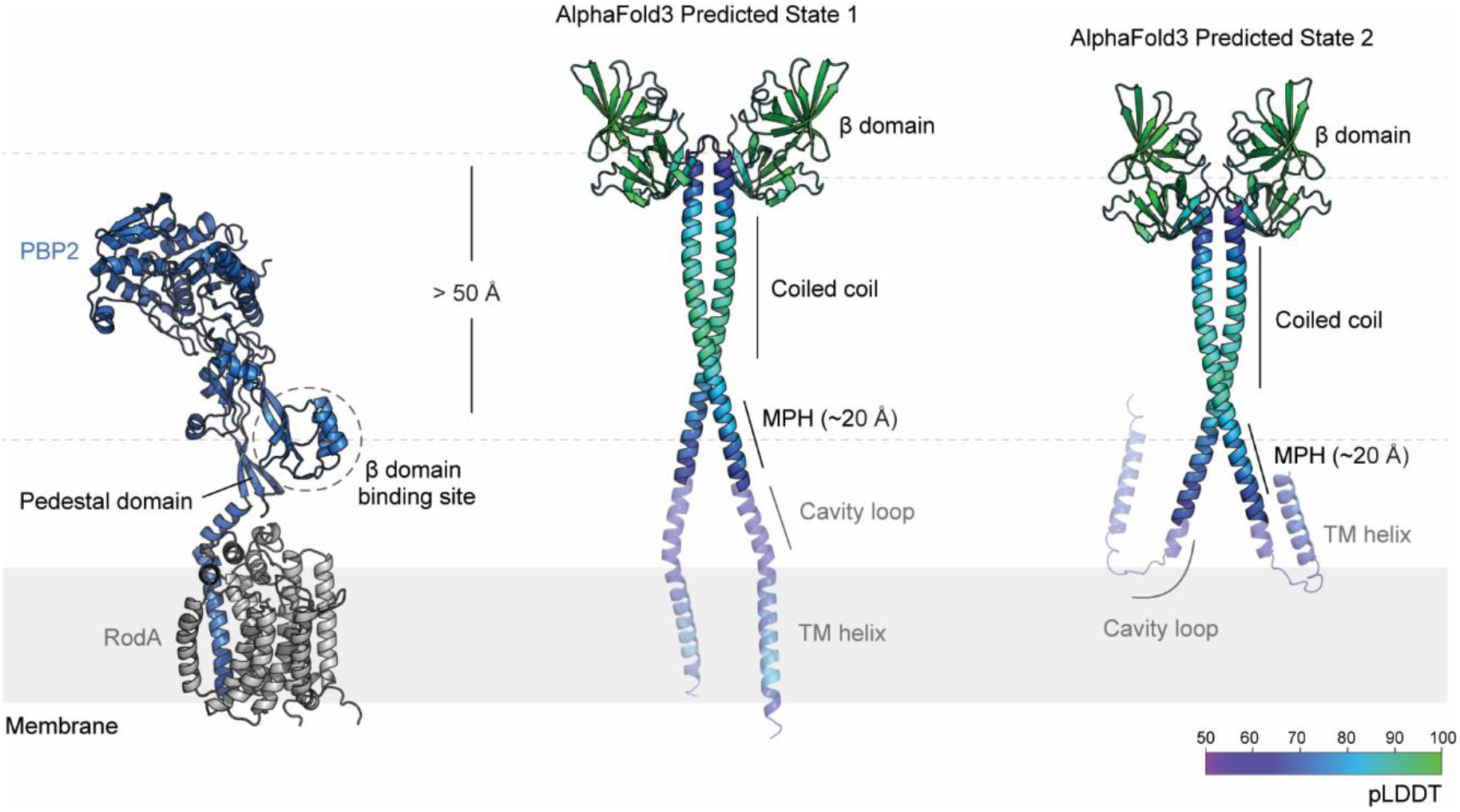
Predictions of the apo-MreC structure are not compatible with PBP2 binding. The AlphaFold3 prediction of the *Tt*RodA-PBP2 complex in the open conformation is shown as ribbons for reference. *Tt*RodA is colored grey and *Tt*PBP2 is blue. The approximate MreC binding site is outlined with a dashed circle. An AlphaFold3 prediction of the dimeric *Tt*MreC structure is shown as ribbons colored according to the pLDDT score. The dashed lines parallel to the membrane indicate the position of the β domain of MreC in the apo state and its binding site on PBP2. The membrane proximal helix adopts a continuous alpha helix with the coiled coil domain. Low confidence regions are shown as transparent cartoons.

**Supplemental Fig. 5.**
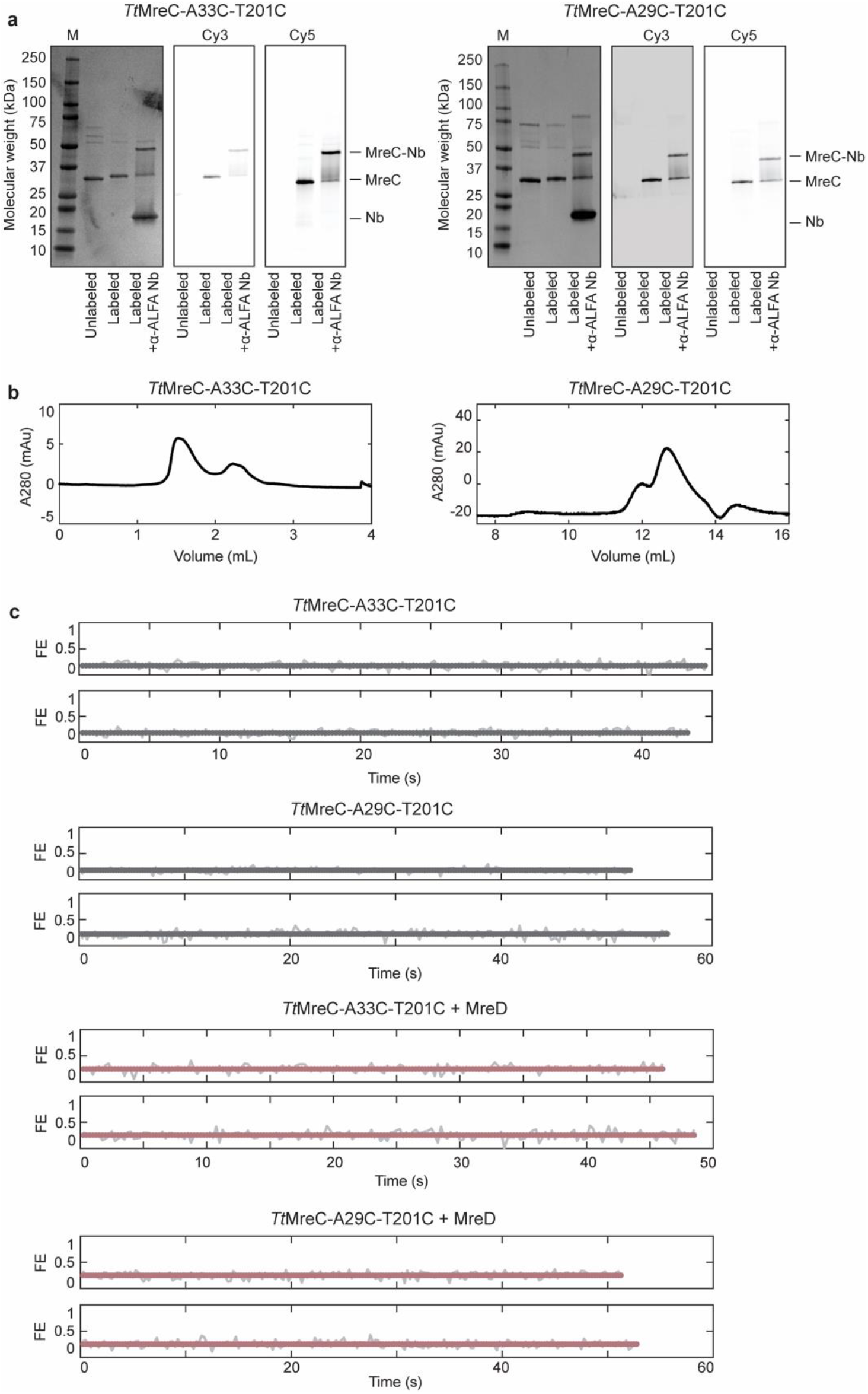
*Tt*MreC smFRET constructs. **a**, Coomassie and fluorescent gels of *Tt*MreC double-cysteine constructs show efficient labeling with both Cy3 and Cy5 dyes and SDS-PAGE stable binding to the α-ALFA nanobody. **b**, Size-exclusion elution profiles of constructs in (**a**) run on 3.2/300 S200I and 10/300 S200I columns, respectively. **c**, Example single-molecule trajectories corresponding to population histograms in Fig. 2c.

**Supplemental Fig. 6.**
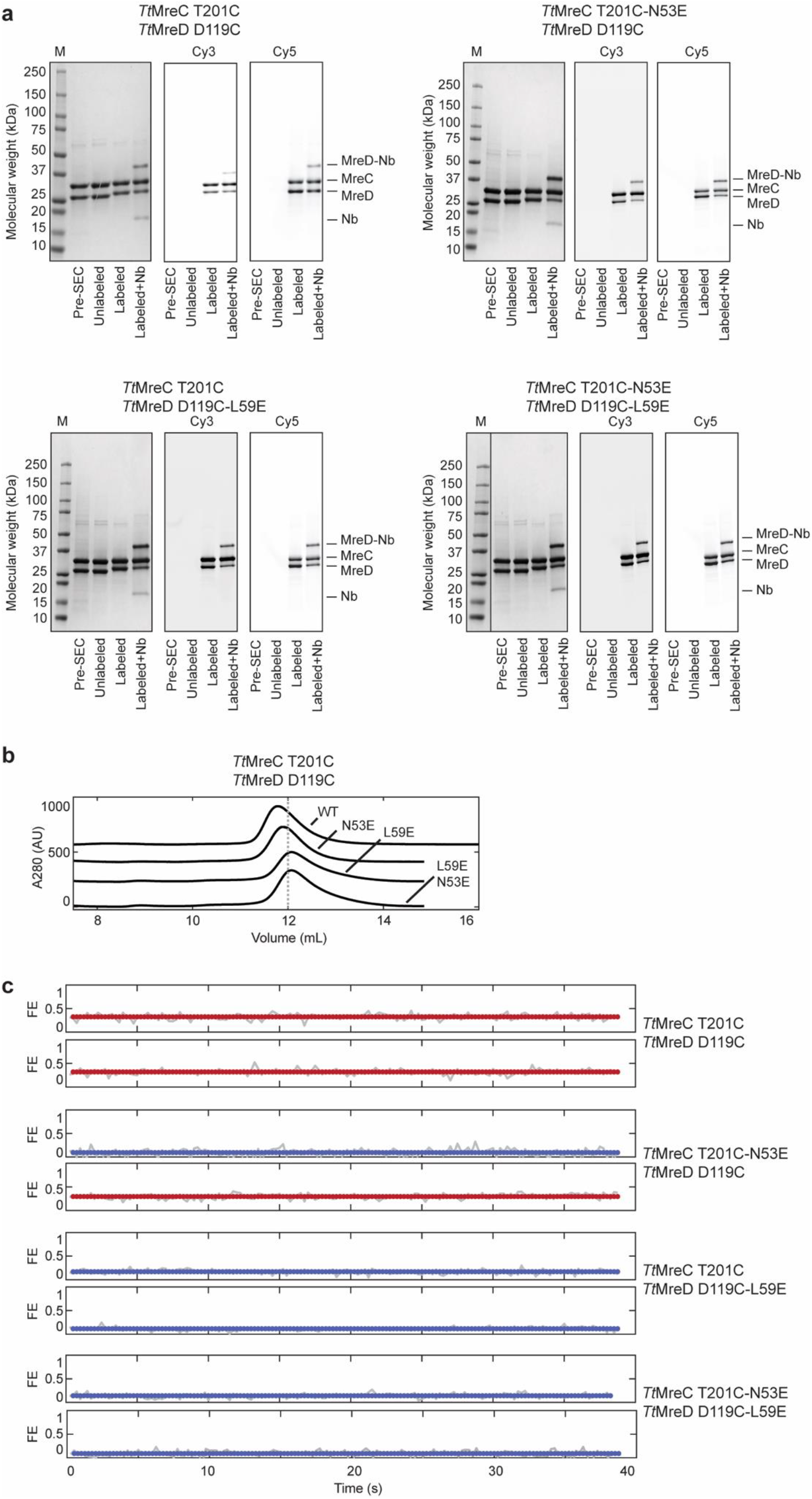
MPH smFRET constructs. **a**, Coomassie and fluorescent gels of *Tt*MreC^T201C^-MreD^D119C^ and MPH mutants show efficient labeling with both Cy3 and Cy5 dyes and SDS-PAGE stable binding to the α-ALFA nanobody. **b**, SEC elution profiles of MPH constructs in (**a**). All variants elute as a monomeric complex of *Tt*MreC-MreD. **c**, Example single-molecule trajectories of *Tt*MreC^T201C^-MreD^D119C^ and MPH mutants corresponding to population histograms in Fig. 3c.

**Supplemental Fig. 7.**
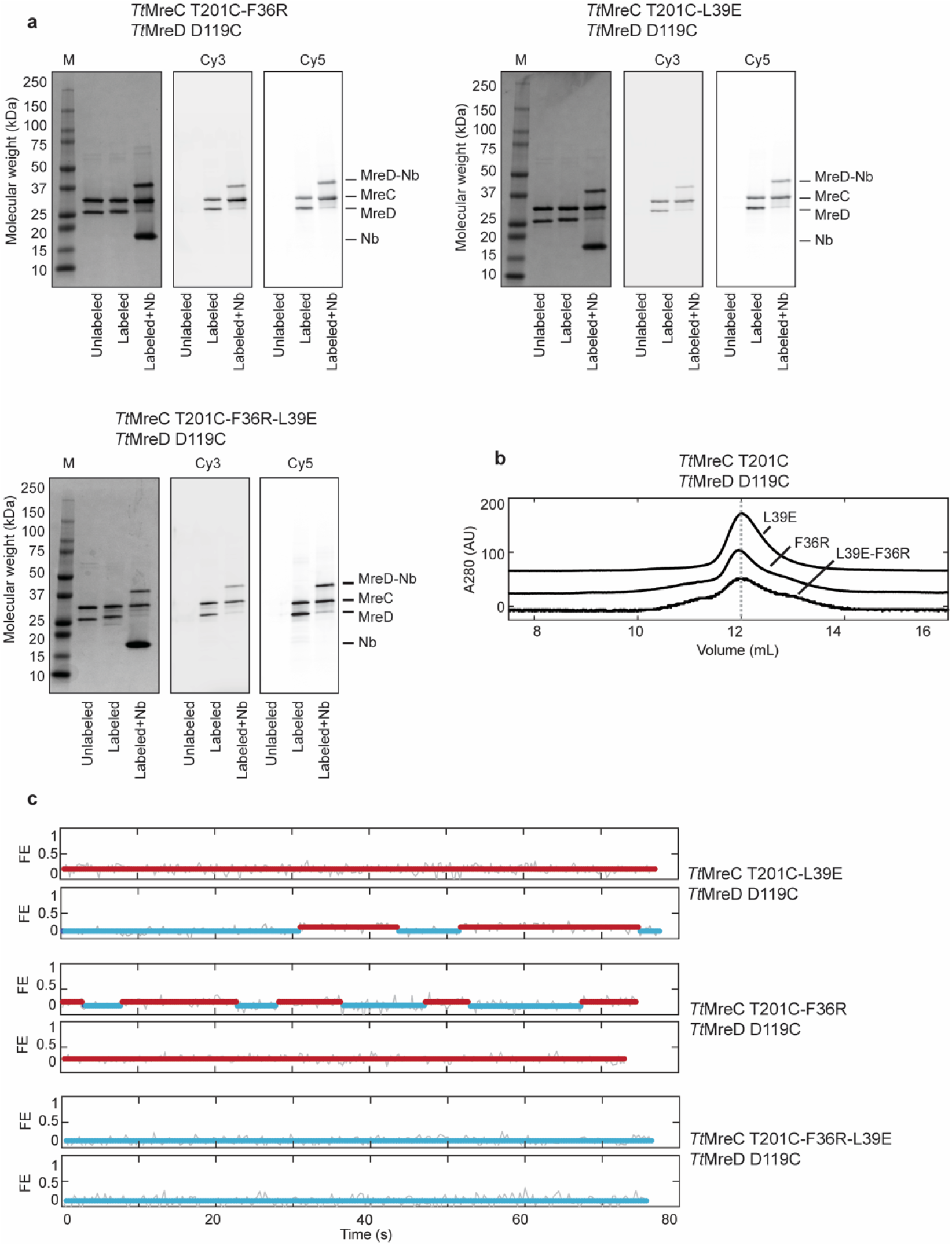
Cavity loop smFRET constructs. **a**, Coomassie and fluorescent gels of *Tt*MreC^T201C^-MreD^D119C^ cavity loop mutants show efficient labeling with both Cy3 and Cy5 dyes and SDS-PAGE stable binding to the α-ALFA nanobody. **b**, SEC elution profiles of cavity loop constructs in (**a**). All variants elute as a monomeric complex of *Tt*MreC-MreD. **c**, Example single-molecule trajectories of cavity loop mutants corresponding to population histograms in Fig. 3c.

**Supplemental Fig. 8.**
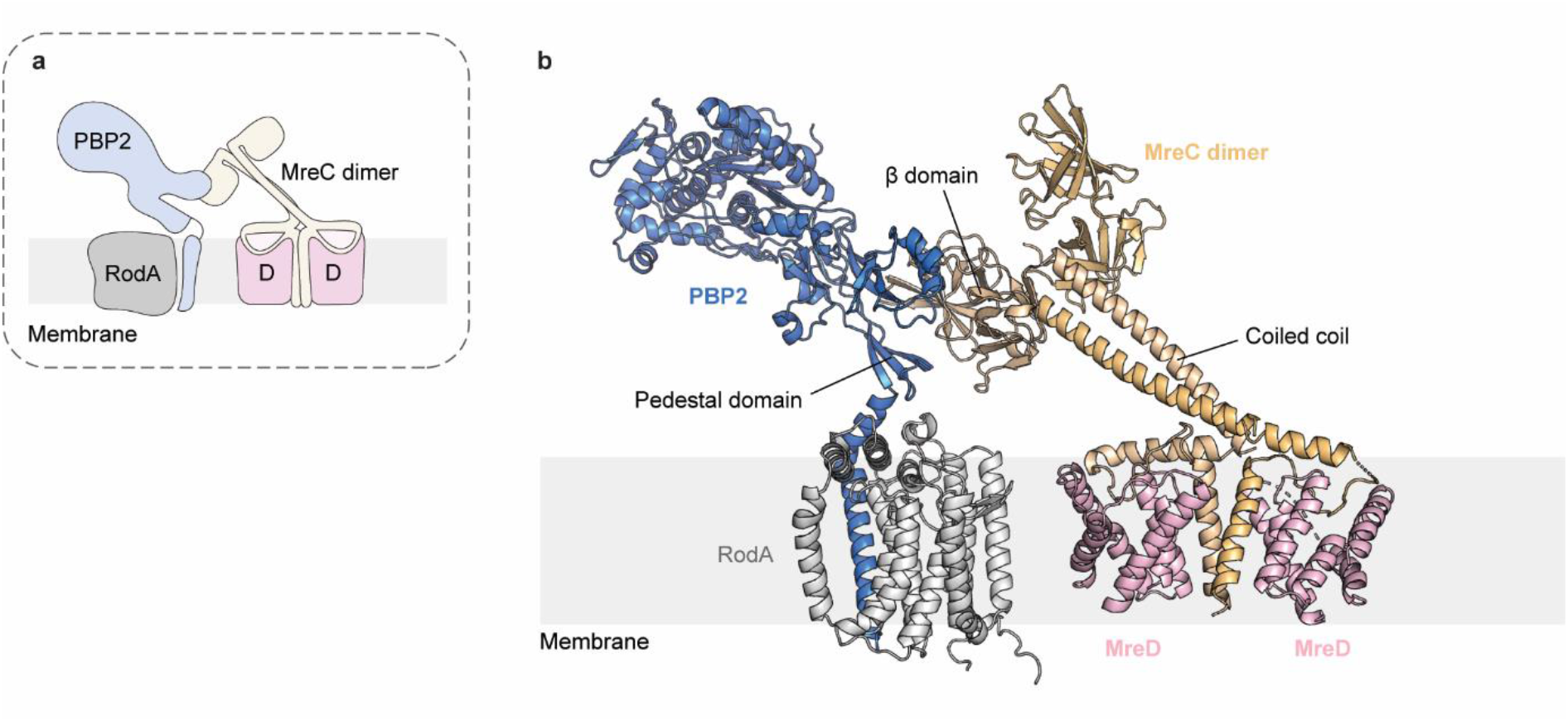
Model of the four-component Rod complex structure. **a**, Cartoon schematic illustrates the predicted structure of the four-component complex. **b**, Ribbon diagram shows the model of the four-component complex generated using AlphaFold3 (AF3) and our cryo-EM structure. *Tt*RodA-PBP2 and a copy of the *Tt*MreC β domain predicted by AF3 was used as a starting complex. The *Tt*MreC coiled coil domain with a single copy of the β domain was isolated from the AF3 prediction. We manually positioned this subcomplex into the experimentally determined cryo-EM map at the maximal degree of tilt. The remaining *Tt*RodA-PBP2-MreC β domain model was aligned on the β domain on the top of the coiled coil. The position of *Tt*RodA-PBP2 was adjusted to impose an approximately planar orientation of the transmembrane domains of all four components by assuming a flexible linkage between the coiled coil and β domain of *Tt*MreC.

**Supplemental Fig. 9.**
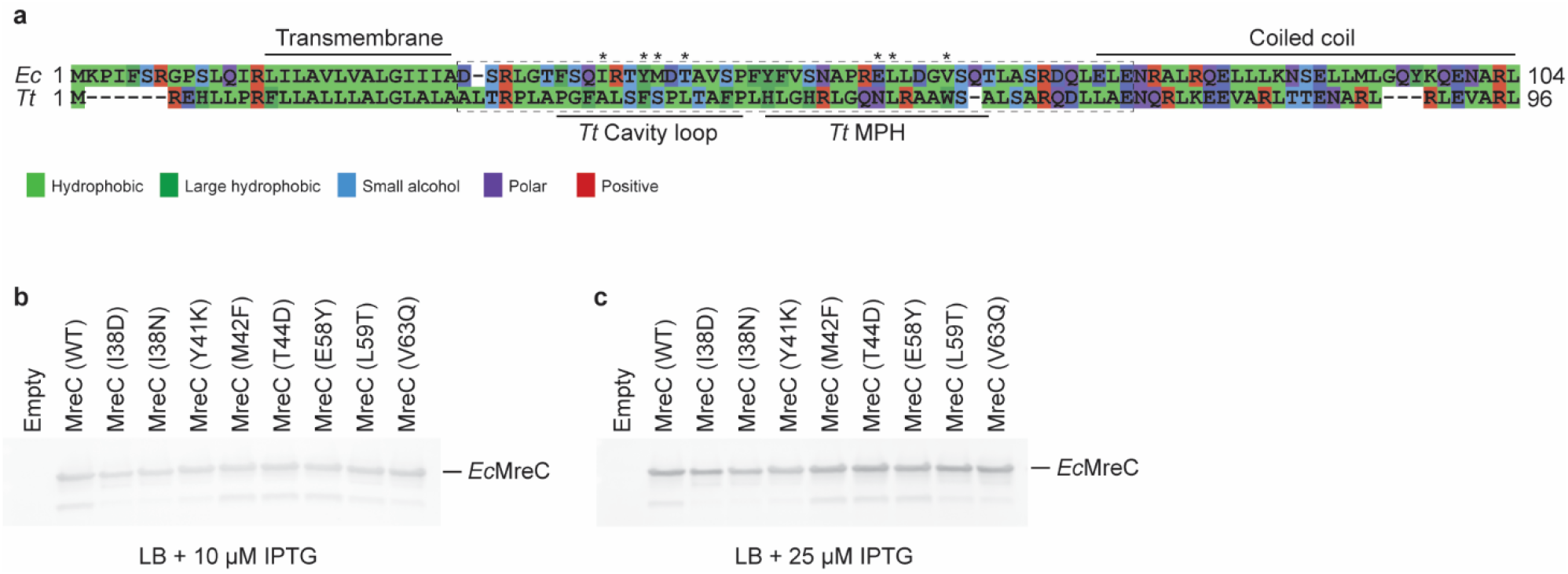
The membrane proximal regions of *Ec*MreC are important for Rod complex function *in vivo*. **a**, The alignment of *Ec* and *Tt*MreC membrane proximal regions was visualized using alignmentviewer.org and colored according to amino acid property. The amino acid sequence is for each protein is overlaid. Dashed box indicates the regions of *Ec*MreC that were included in the site-saturation mutagenesis library. Asterisks indicate mutants that were isolated from the library screen. Western blot analysis of the strains analyzed in the *ΔmreC* complementation assay demonstrating expression of all variants of *Ec*MreC at **b**, 10 μM and **c**, 25 μM IPTG.

**Supplemental Fig. 10.**
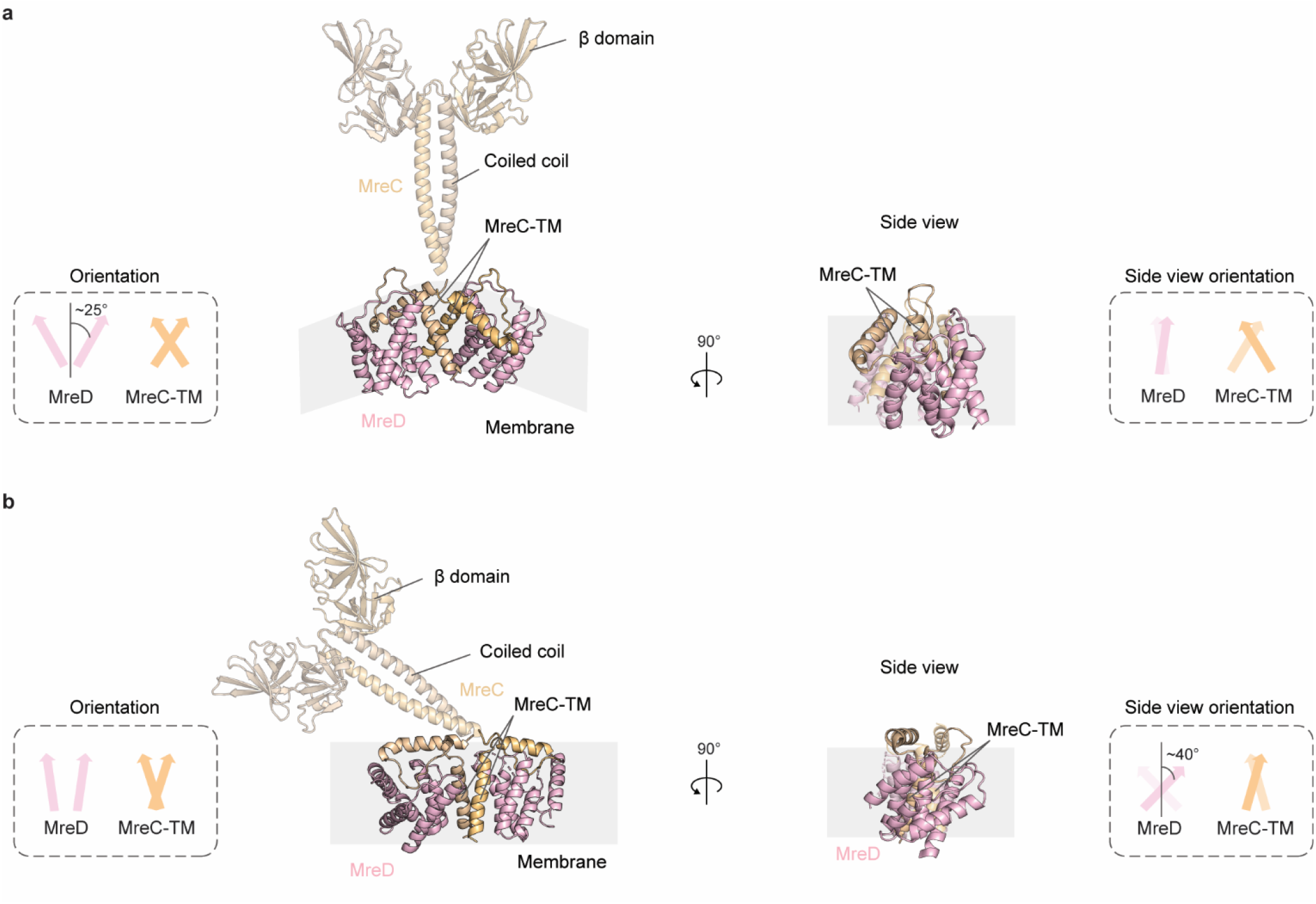
*Tt*MreD adopts two tilted orientations. **a**, The *Tt*MreC-MreD-BRIL (flexible C-terminal fusion) + α-BRIL Fab structure is shown as ribbons, colored as in Fig. 1a (left). The two β domains and the coiled coil of *Tt*MreC are shown as tan ribbons depicting an average of the possible orientations of this region. The approximate position of the membrane is annotated in grey. A 90° rotation about the axis perpendicular to the membrane is shown without the coiled coil and β domains (right). The insets depict the orientations of each copy of *Tt*MreD and the transmembrane helices of *Tt*MreC. In the lower resolution structure, the *Tt*MreD proteins are tilted by approximately 25°. **b**, The same panels as in (**a**) but for the 3.6 Å resolution structure of *Tt*MreC-MreD with the rigid internal fusion of BRIL. In this structure, the *Tt*MreD proteins are tilted by approximately 40° when viewed from the side.

**Supplemental Table 1.** Cryo-EM data collection, refinement and validation statistics.

| | *Tt*MreC-MreD-BRIL internal<br>fusion + α-BRIL Fab in LMNG<br>(PDB: 9DVB)<br>(EMD-47199) | *Tt*MreC-MreD-BRIL flexible<br>fusion + α-BRIL Fab in DDM<br>(PDB: 9DVC)<br>(EMD-47200) |
| --- | --- | --- |
| **Data collection and processing** |  |  |
| Microscope | Talos Arctica | Talos Arctica |
| Magnification | 36,000 | 36,000 |
| Voltage (kV) | 200 | 200 |
| Electron exposure (e-/Å<sup>2</sup>) | 52 | 61 |
| Defocus range (μm) | ~1.0-3.0 | ~0.8-2.5 |
| Pixel size (Å) | 1.1 | 1.1 |
| Symmetry imposed | C1 | C1 |
| Initial particle images (no.) | 2,973,348 | 1,285,219 |
| Final particle images (no.) | 166,346 | 51,857 |
| Map resolution (Å) | 3.6 | 7.4 |
| FSC threshold | 0.143 | 0.143 |
| **Refinement** |  |  |
| Initial model used<br>(PDB code) | *Tt*MreC-MreD-BRIL<br>(AlphaFold3)<br>α-BRIL Fab (7C61) | 9DVB<br>(Rigid body refinement only) |
| Model composition |  |  |
| Non-H atoms | 10,802 | 6,838 |
| Protein residues | 1,410 | 889 |
| Ligands | 0 | 0 |
| Model vs. data CC |  |  |
| Overall | 0.82 | 0.68 |
| For ligands | n/a | n/a |
| R.m.s. deviations |  |  |
| Bond lengths (Å) | 0.004 | 0.005 |
| Bond angles (°) | 0.911 | 0.926 |
| Validation |  |  |
| MolProbity score | 1.27 | 1.71 |
| Clashscore | 5.04 | 16.00 |
| Poor rotamers (%) | 0.52 | 0.84 |
| Ramachandran plot |  |  |
| Favored (%) | 98.32 | 98.62 |
| Allowed (%) | 1.68 | 1.38 |
| Disallowed (%) | 0 | 0 |

**Supplemental Table 2.**
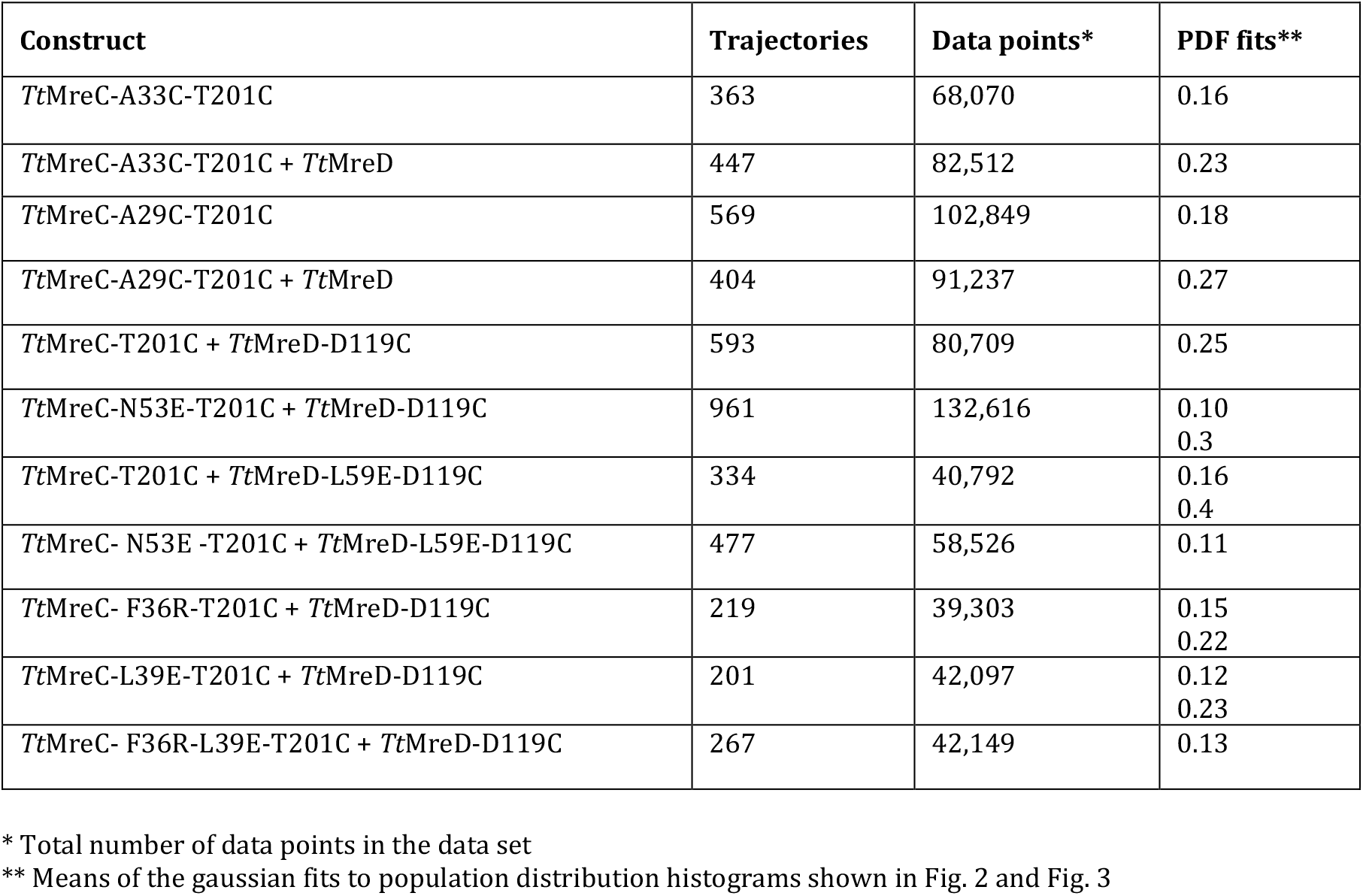
smFRET analysis and fitting.

| **Construct** | **Trajectories** | **Data points\*** | **PDF fits\*\*** |
| --- | --- | --- | --- |
| *TtMreC*-A33C-T201C | 363 | 68,070 | 0.16 |
| *TtMreC*-A33C-T201C + *TtMreD* | 447 | 82,512 | 0.23 |
| *TtMreC*-A29C-T201C | 569 | 102,849 | 0.18 |
| *TtMreC*-A29C-T201C + *TtMreD* | 404 | 91,237 | 0.27 |
| *TtMreC*-T201C + *TtMreD*-D119C | 593 | 80,709 | 0.25 |
| *TtMreC*-N53E-T201C + *TtMreD*-D119C | 961 | 132,616 | 0.10<br>0.3 |
| *TtMreC*-T201C + *TtMreD*-L59E-D119C | 334 | 40,792 | 0.16<br>0.4 |
| *TtMreC*- N53E -T201C + *TtMreD*-L59E-D119C | 477 | 58,526 | 0.11 |
| *TtMreC*- F36R-T201C + *TtMreD*-D119C | 219 | 39,303 | 0.15<br>0.22 |
| *TtMreC*-L39E-T201C + *TtMreD*-D119C | 201 | 42,097 | 0.12<br>0.23 |
| *TtMreC*- F36R-L39E-T201C + *TtMreD*-D119C | 267 | 42,149 | 0.13 |

\* Total number of data points in the data set
\*\* Means of the gaussian fits to population distribution histograms shown in Fig. 2 and Fig. 3

**Supplemental Table 3.** Library screening results.

| Clone ID | Substitution | Region | Gamma domain mutation |
| --- | --- | --- | --- |
| 1 | G33T | Loop | R292H |
| 2 | E58Y | MPH | none |
| 3 | T44D | Loop | none |
| 4 | Y41K | Loop | none |
| 5 | I38N | Loop | none |
| 6 | L73P | MPH | E116K |
| 7 | M42F | Loop | none |
|  | V63Q | MPH | none |
| 8 | G62F | MPH | E116K |
| 9 | I38D | Loop | none |
| 10 | L59T | MPH | none |
| 11 | L32V | Loop | E116D |
| 12 | V46C | Loop | none |

**Supplemental Table 4.** *In vitro* plasmids.

| Plasmid | Genotype | Source |
| --- | --- | --- |
| pMG7 | *pCDF-Duet-TtMreC-3C-PrtC* | This study |
| pMG36 | *pTarget α-BRIL Fab Heavy chain CD\** | Mukherjee *et al.*, 2020<br>Skiba *et al.*, 2024<br>Bailey *et al.*, 2018 |
| pMG113 | *pCDF-Duet-TtMreC-S37C-3C-PrtC* | This study |
| pMG114 | *pCDF-Duet-TtMreC-T40C-3C-PrtC* | This study |
| pMG116 | *pET-Duet-MVKIH-SUMO-FLAG-3C-TtMreD-BRIL-Q17C* | This study |
| pMG117 | *pET-Duet-MVKIH-SUMO-FLAG-3C-TtMreD-BRIL-E22C* | This study |
| pMG126 | *pCDF-Duet-TtMreC-TM-GGS-PrtC* | This study |
| pMG127 | *pCDF-Duet-TtMreC-TM-CL-GGS-PrtC* | This study |
| pMG128 | *pCDF-Duet-TtMreC-TM-CL-MPH-GGS-PrtC* | This study |
| pMG54 | *pET-Duet-SUMO-FLAG-3C-TtMreD-BRIL* | This study |
| pSI214 | *pCDF-Duet-PrtC-SwapCC-MreC-T201C* | This study |
| pSI291 | *pCDF-Duet-PrtC-ALFA-SwapCC-MreC-A33C-T201C* | This study |
| pSI292 | *pCDF-Duet-PrtC-ALFA-SwapCC-MreC-A29C-T201C* | This study |
| pSI324 | *pCDF-Duet-PrtC-SwapCC-MreC-F36R-T201C* | This study |
| pSI325 | *pCDF-Duet-PrtC-SwapCC-MreC-L39E-T201C* | This study |
| pSI329 | *pCDF-Duet-PrtC-SwapCC-MreC-F36R-L39E-T201C* | This study |
| pDSG7 | *pET-Duet-SUMO-FLAG-3C-TtMreD-internal-BRIL* | This study |
| pDSG18 | *pCDF-Duet-PrtC-SwapCC-MreC-N53E-T201C* | This study |
| pDSG22 | *pET-Duet-FLAG-ALFA-TtMreD-D119C-BRIL* | This study |
| pDSG26 | *pET-Duet-FLAG-ALFA-TtMreD-L59E-D119C-BRIL* | This study |
| pMAS266 | *pD2610 α-BRIL Fab light chain* | Mukherjee *et al.*, 2020<br>Skiba *et al.*, 2024 |

**Supplemental Table 5.**
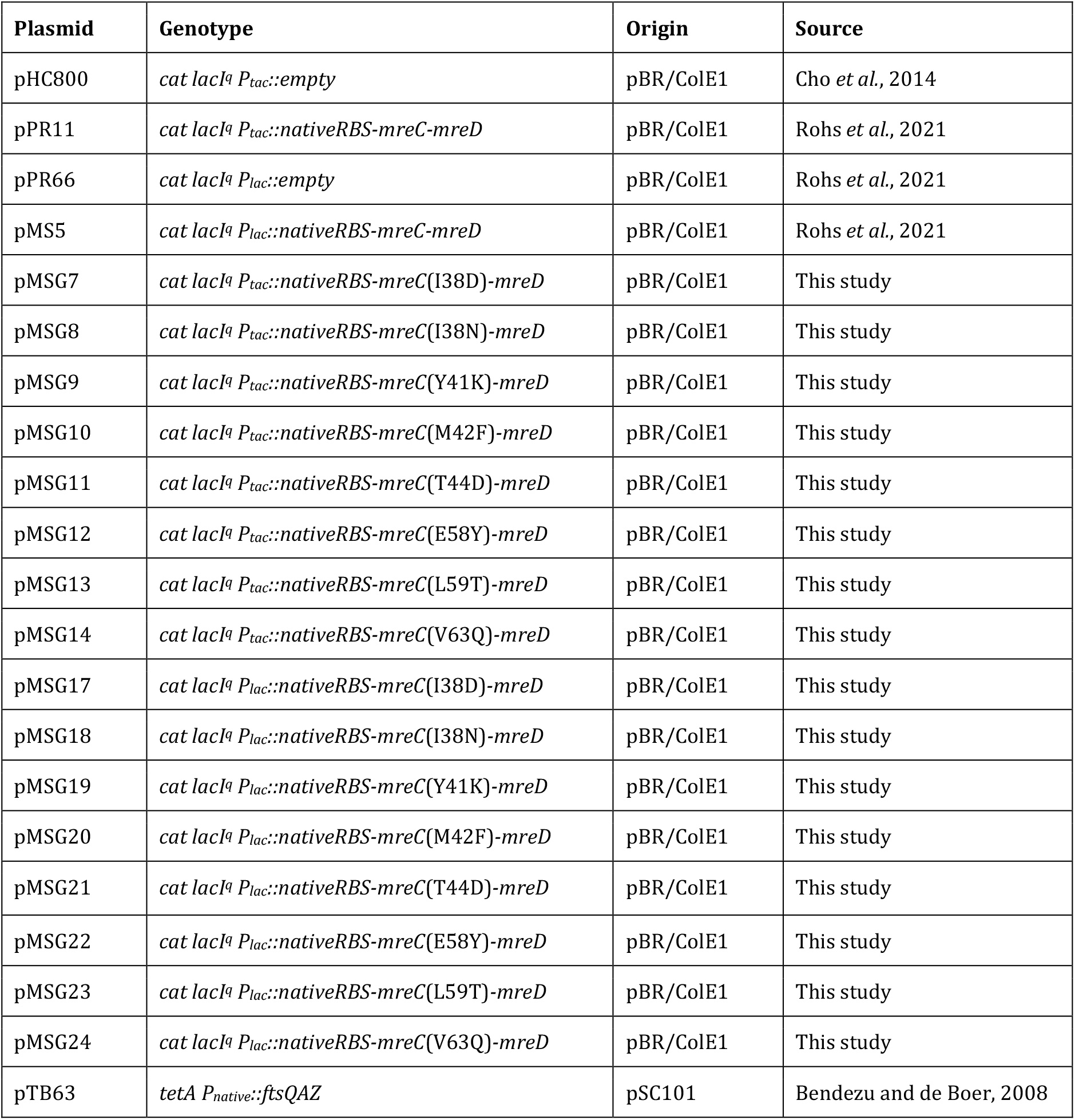
*In vivo* plasmids.

